# Neural tube closure requires the endocytic receptor Lrp2 and its functional interaction with intracellular scaffolds

**DOI:** 10.1101/2020.07.15.205252

**Authors:** Izabela Kowalczyk, Chanjae Lee, Elisabeth Schuster, Josefine Hoeren, Valentina Trivigno, Levin Riedel, Jessica Görne, John B. Wallingford, Annette Hammes, Kerstin Feistel

## Abstract

Recent studies have revealed that pathogenic mutations in the endocytic receptor LRP2 in humans are associated with severe neural tube closure defects (NTDs) such as anencephaly and spina bifida. Here, we combined analysis of neural tube closure in mouse and in the African Clawed Frog *Xenopus laevis* to elucidate the etiology of Lrp2-related NTDs. *Lrp2* loss-of-function (LOF) impaired neuroepithelial morphogenesis, culminating in NTDs that impeded anterior neural plate folding and neural tube closure in both model organisms. Loss of Lrp2 severely affected apical constriction as well as proper localization of the core planar cell polarity (PCP) protein Vangl2, demonstrating a highly conserved role of the receptor in these processes essential for neural tube formation. In addition, we identified a novel functional interaction of Lrp2 with the intracellular adaptor proteins Shroom3 and Gipc1 in the developing forebrain. Our data suggest that during neurulation, motifs within the intracellular domain of Lrp2 function as a hub that orchestrates endocytic membrane removal for efficient apical constriction as well as PCP component trafficking in a temporospatial manner.

**Summary statement:** Analysis of neurulation in mouse and *Xenopus* reveals novel roles for Lrp2-mediated endocytosis in orchestrating apical constriction and planar cell polarity essential for neural tube closure.

## Introduction

Morphogenesis of the forebrain at the anterior end of the neural tube is a complex process. In higher vertebrates, the forebrain originates from a simple sheet of neuroepithelial cells and subsequently forms the largest part of the brain. The anterior neural plate (NP) evaginates, bends, and then progressively fuses along the dorsal midline to establish the neural tube (Nikolopoulou et al., 2017). Defects in these pivotal processes during early brain development lead to a wide range of congenital brain malformations in humans, including holoprosencephaly (HPE) and anencephaly. Several environmental and genetic risk factors have been identified as possible causes of structural brain anomalies (Greene and Copp, 2014; Wallingford et al., 2013).

The low-density lipoprotein (LDL) receptor-related protein 2 (LRP2, also known as megalin; Saito et al., 1994) is a key gene associated with severe forebrain defects. LRP2 is a multifunctional cell surface receptor that is structurally related to the LDL receptor family (Nykjaer and Willnow, 2002) and localizes to the apical surface of epithelia. All LRP2 orthologs share a large extracellular and a comparatively short intracellular domain that harbors not only typical NPxY endocytosis motifs (Chen et al., 1990) but also phosphorylation and PDZ-binding motifs relevant for interaction with intracellular adaptors and scaffolding proteins (Gotthardt et al., 2000; Naccache et al., 2006). Humans with autosomal recessive *LRP2* gene defects suffer from Donnai-Barrow syndrome (DBS) with a range of clinical characteristics including craniofacial anomalies (ocular hypertelorism, enlarged fontanelle) and forebrain defects such as agenesis of the corpus callosum (Kantarci et al., 2007; Ozdemir et al., 2020) and microforms of HPE (Rosenfeld et al., 2010). Severe forms of HPE in families presenting with oligogenic events involving *LRP2* have also been described (Kim et al., 2019). Most of the clinical characteristics seen in patients with *LRP2* gene mutations are also present in the LRP2-deficient mouse (Cases et al., 2015; Hammes et al., 2005; Kur et al., 2014; Spoelgen et al., 2005; Wicher and Aldskogius, 2008), making it a valuable model for studying the mechanistic origin of this human congenital disorder.

We previously identified LRP2 as a novel component of the sonic hedgehog (SHH) signaling machinery in the ventral midline of the developing forebrain (Christ et al., 2012). Interestingly, LRP2-deficient mice not only suffer from HPE caused by ventral forebrain defects, but also display defects of the anterior dorsolateral neural tube as well as spinal cord anomalies that cannot be explained by loss of SHH signaling in the developing forebrain (Kur et al., 2014; Wicher and Aldskogius, 2008; Ybot-Gonzalez et al., 2002). In previous studies, we and other labs demonstrated that LRP2 null mutants present with a dilated dorsal neural tube and in 38 % of all mice we observed cranial neural tube closure defects (NTDs; Kur et al., 2014; Sabatino et al., 2017). Of note, human *LRP2* variants have now also been identified in patients suffering from NTDs, ultimately leading to anencephaly and myelomeningocele (open spina bifida; Rebekah Prasoona et al., 2018; Renard et al., 2019). Neural tube closure (NTC) and morphogenesis is marked by extensive and rapid cell and tissue rearrangements, driven by morphogenetic events such as cell migration and intercalation as well as cell shape changes (Wallingford, 2005). Since LRP2 is a candidate gene for NTDs, we asked whether Lrp2-mediated endocytosis was required for dynamic cell behaviors during NTC.

Here, we used the African Clawed Frog *Xenopus laevis* and the mouse, showing for the first time that there are common NTDs in Lrp2-deficient *Xenopus* and mouse embryos. Loss of LRP2-mediated endocytosis impaired apical constriction and caused aberrant localization of the core planar cell polarity (PCP) component Vangl2 in both model organisms. Lrp2 functionally interacted with intracellular adaptor and scaffold proteins to exert its function in NTC.

## Results

### Constricting neuroepithelial cells in the forebrain show enriched localization of LRP2 on the apical surface

To evaluate the feasibility of using *Xenopus laevis* to study the etiology of LRP2-related NTDs, we first analyzed *lrp2* expression in relevant stages. Lrp2 protein is maternally expressed and protein levels increase with the onset of zygotic *lrp2* transcription (Fig. S1A; Peshkin et al., 2019; Session et al., 2016). *lrp2* was also expressed in DBS-relevant organ anlagen such as the brain, eye, otic vesicle and pronephros (Fig. S1B).

Neuroepithelial cells undergo complex shape changes to apically constrict and form hinge points that ultimately allow proper tissue morphogenesis during neural tube upfolding. *lrp2* transcripts were detected in the NP (Fig. 1A) and the protein localized in *Xenopus* (Fig. 1B) and mouse (Fig. 1C, C’, C”) neuroepithelial cells during early forebrain development. Interestingly, the most prominent signals for Lrp2 were detected in apically constricting cells, which in *Xenopus* are first apparent along the borders of the NP (arrowheads in Fig. 1B; B’) and appear upon formation of the optic evagination (OE) in mouse (Fig. 1C’). Stimulated Emission Depletion (STED) microscopy further revealed LRP2 localization concentrated in the periciliary region (Fig. 1D), a highly endocytic plasma membrane domain at the base of the primary cilium (Benmerah, 2013; Molla-Herman et al., 2010).

**Fig. 1:**
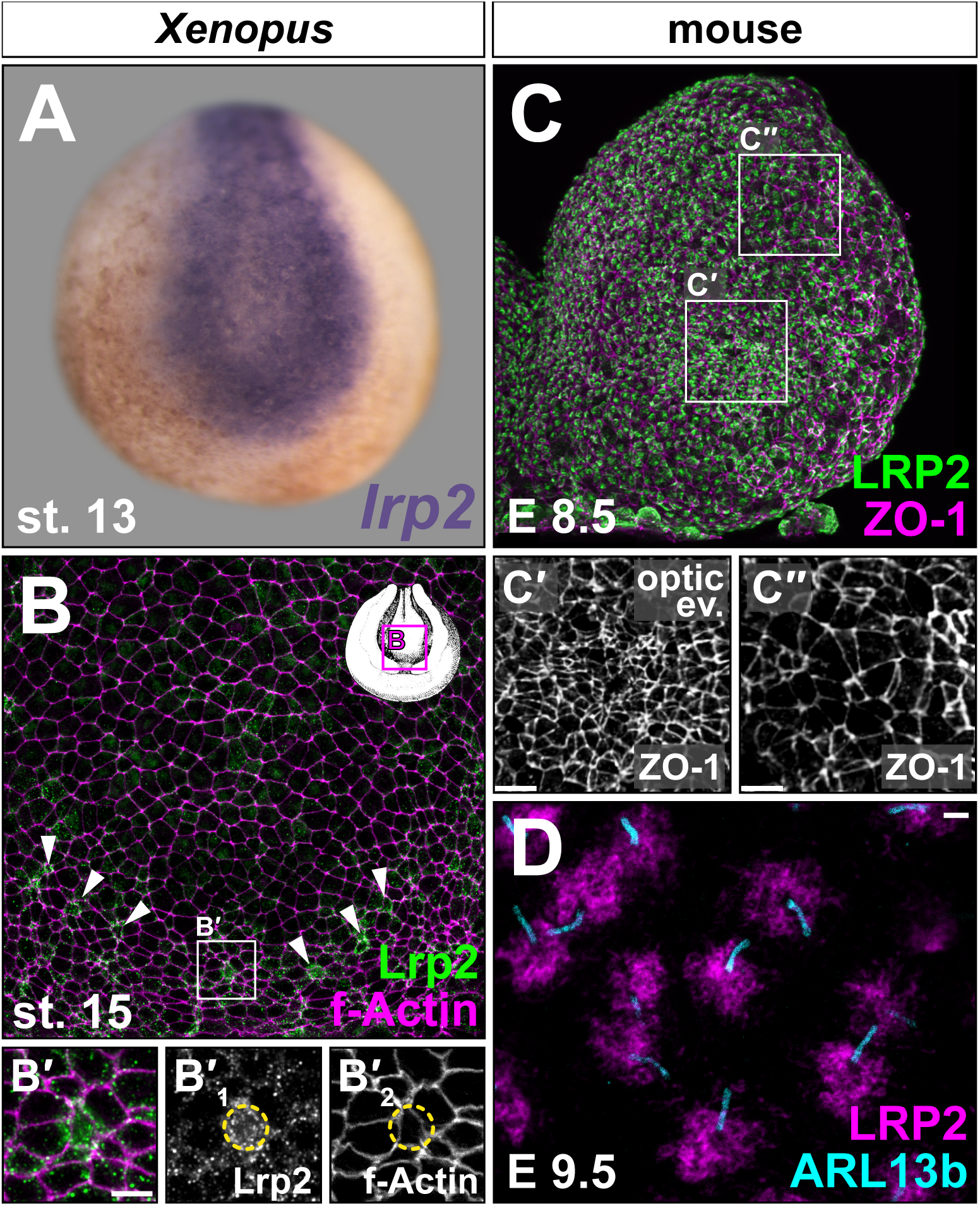
Lrp2 is expressed in the neuroepithelium and increased in constricting cells. *lrp2* mRNA (**A**) and protein (**B-D**) expression analyzed by *in situ* hybridization and immunofluorescence, respectively. Neurula stage (st.) embryos (frontal en face views, dorsal up). (**A**) Neural *lrp2* expression at st. 13. (**B**) St. 15 forebrain region; Lrp2 present in most cells (outlined by f-Actin); single cells with higher Lrp2 levels localized along anterior rim of neural folds (NFs; arrowheads). (**B’**) Magnification as indicated in (**B**); increased Lrp2 levels in cells with small apical surface (circle on single channels **B’**_1_ and **B’**_2_). (**C**) LRP2 detected throughout embryonic day (E) 8.5 anterior NFs (left hemisphere shown), concentrated in areas undergoing apical constriction (compare (**C’**) – apically constricted cells in optic evagination – and (**C”**) – dorsolateral cells with larger cell surface). Cell boundaries marked by tight junction protein ZO-1. (**D**) STED imaging; condensation of LRP2 around primary cilia (ARL13b^+^) of neuroepithelial cells at E9.5. Scale bars (**B’, C’, C”**): 10 μm; (**D**): 1 μm.

The highly specific localization of Lrp2 / LRP2 in neuroepithelial cells, which clearly correlated with cell shape changes during the process of apical constriction (AC), suggested a role for this receptor in cellular remodeling to enable tissue morphogenesis.

### Loss of Lrp2 disrupts neural tube morphogenesis

We next examined neurulation in *Xenopus* upon *lrp2* loss-of-function (LOF) and in mouse null mutants for *Lrp2*. Genetically modified mouse *Lrp2^−/−^* embryos presented with altered neural tube morphology at embryonic day (E) 8.5 compared to wild type (WT) controls, as observed using scanning electron microscopy (Fig. 2A-F). Following the neural tube morphogenetic processes in WT and somite-matched mutant embryos from 6 to 8-somite stages, we detected a delay and ultimately a deficit in OE formation in the developing forebrain of *Lrp2^−/−^* mice (Fig. 2D-F). Neural fold morphogenesis was also impaired in *Lrp2* mutant embryos during neurulation, which became especially evident at the 8-somite stage, when mutant neural folds were less elevated (Fig. 2F) compared to controls (Fig. 2C). In WT embryos at E9.5, the anterior neural tube is closed and midline separation of the forebrain vesicles starts. Compared to the WT (Fig. S2A, C), *Lrp2* null mutants at this stage had either severely dilated (65%, 22 / 34; Fig. S2B) or open neural tubes (35%, 12 / 34; Fig. S2D). At E18.5, when a proper skin-covered skull had formed in WT embryos (Fig. S2E), we observed exencephaly and anencephaly in *Lrp2* null mutants (Fig. S2F), presumably a consequence of failed neural tube closure.

**Fig. 2:**
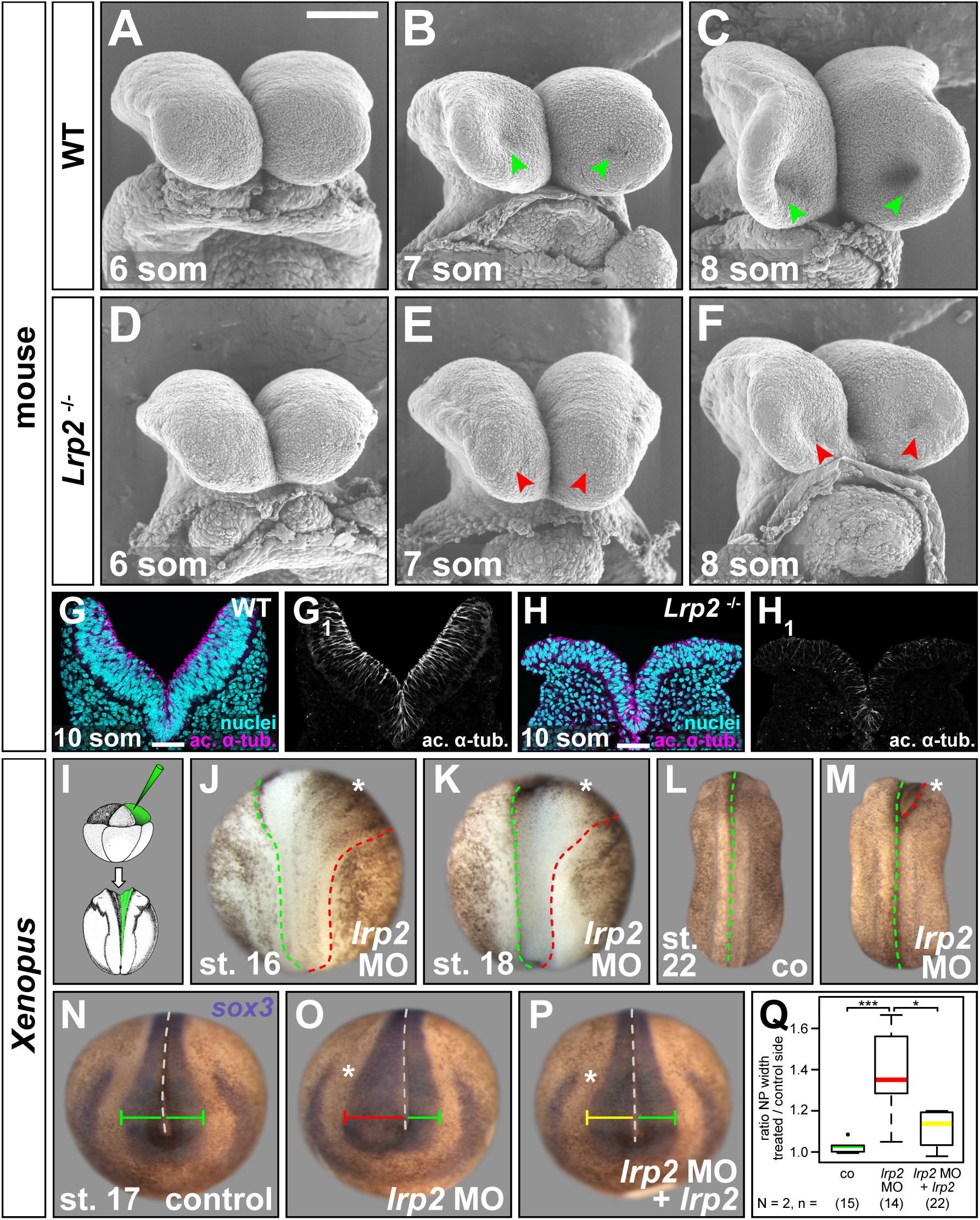
Lrp2 is required for proper neuroepithelial morphogenesis and neural tube closure. Analysis of neural plate (NP) morphology. (**A-F**) Scanning electron microscopy images of neural folds (NFs) from embryonic day (E) 8.5 wild type (WT) and *Lrp2^−/−^* mouse embryos at 6, 7 and 8-somite (som) stage (st.), frontal views. (**A-C**) WT NFs progressively elevated, optic evagination initiated (arrowheads in **B, C**). (**D-F**) Narrower NFs, delayed elevation, impaired optic evagination in mutants (arrowheads in **E, F**). (**G, H**) Immunofluorescence staining detecting acetylated α-tubulin (ac. α-tub.) and DAPI-stained nuclei on coronal sections of 10-somite WT and *Lrp2^−/−^* mouse NF. Scale bars (**A-F**): 100 μm; (**G, H**): 50 μm. (**I**) 8-cell embryos injected into one dorsal animal blastomere to target the NP unilaterally. (**J, K**) Morpholino oligomer targeting *lrp2*.L (*lrp2* MO) impaired hingepoint formation and convergent extension on injected side (asterisk) during NF elevation between st. 16 (**J**) and st. 18 (**K**). (**L, M**) After neural tubes closed in controls (co; **L**), anterior NFs on injected side (asterisk in **M**) remained open. (**N-Q**) *In situ* hybridization using *sox3* probe revealed normal NP width (green bar) in co (**N**); *lrp2* MO-impaired NP narrowing (red bar, **O**) partially rescued (yellow bar) by re-introduction of *lrp2* (**P**). (**Q**) Graphical representation of results from (**N-P**); Wilcoxon rank sum test.

Integrity of the neuroepithelium was further evaluated in coronal sections of anterior neural tissue at E8.5. Neural folds of 10-somite WT embryos were elevated and staining for acetylated α-tubulin showed the distribution of stabilized tubulin, highlighting the apicobasal axis of the pseudostratified neuroepithelial sheet (Fig. 2G, G_1_). In somite-matched *Lrp2* null mutants, neural fold elevation was impaired and the profound defects in the appearance of the neuroepithelium (Fig. 2H, H_1_) suggested impaired apicobasal elongation, a common feature in cells with defects in AC.

We next looked for a conserved function of *lrp2* in *Xenopus*. A translation-blocking Morpholino oligomer (MO) binding within the 5’ UTR of *lrp2*.L was targeted to neural tissue by injecting into the animal pole of dorsal animal blastomeres of 4-8-cell embryos. Unilateral injections were performed such that the uninjected contralateral sides served as an internal control. This procedure led to a reliable loss of Lrp2 protein in targeted cells comparable to Lrp2^−/−^ mice (Fig. S1C, D) and caused defective NP folding in all embryos in which lineage tracer fluorescence confirmed correct targeting of NP cells (Fig. 2I; supp. movie 1). While pigment accumulation along the border of neural / non-neural tissue illustrated the formation of hinge points on the uninjected side (green dashed line in Fig. 2J, K), hinge point formation was missing along the neural / non-neural border on the injected side (red dashed line in Fig. 2J, K). At the same time, failure of the caudal NP to narrow indicated that convergent extension (CE) was impaired on the injected side (Fig. 2J, K). After neural tube closure in controls (Fig. 2L), anterior neural folds on the *lrp2* MO-injected side remained open (Fig. 2M), confirming that loss of LRP2 / Lrp2 in mouse and *Xenopus* led to severe NTDs.

To quantify the neural phenotype caused by Lrp2 deficiency in *Xenopus*, the NP was labeled using the pan-neural marker *sox3*, NP width in the forebrain region was measured on either side of the midline and displayed as ratios between control and injected side (Fig. 2N-Q). While uninjected control NPs had a ratio around 1 (Fig. 2N, Q), *lrp2* LOF significantly impaired NP narrowing (Fig. 2O, Q). Due to its size of 4663 aa, expressing a full-length construct of *Xenopus lrp2* was not feasible, therefore we used a well-characterized extracellularly truncated construct of human *LRP2*, containing the fourth ligand binding domain as well as transmembrane and intracellular domains (Yuseff et al., 2007), which we refer to as *lrp2* in rescue experiments. Reintroducing *lrp2* on the injected side partially rescued this NP defect (Fig. 2P, Q), confirming specificity of the MO approach and supporting a direct function of Lrp2 in neural tube morphogenesis.

Specificity of the *lrp2* MO was further affirmed by CRISPR/Cas9-mediated genome editing of *lrp2*.L (Fig. S2G), as injection of Cas9 ribonucleic particles (CRNPs) assembled with two different single guide (sg) RNAs into zygotes recapitulated the shortened and widened NP of morphants (Fig. S2H-K), a phenotype that was rescued by co-injection of *lrp2* (cf. Fig. 6L). Lrp2 protein reduction (Fig. S2L-O) as well as sequencing of targeted regions (Fig. S2P, Q) confirmed successful *lrp2* LOF upon CRISPR/Cas9 treatment.

### Lrp2 is cell-autonomously required for efficient apical constriction

Hinge point formation leading to NP bending is a process driven by AC, i.e. narrowing of the apical surface and widening of basolateral cell aspects (Martin and Goldstein, 2014). The loss of hinge points in *Xenopus* prompted us to examine whether Lrp2 plays a role in regulating such cell shape changes. To this end we analyzed the morphology of *Xenopus* neuroepithelial cells upon injection of *lrp2* MO (Fig. 3A-F). f-Actin staining, delineating the cell circumference, revealed a much larger apical surface in cells that had received *lrp2* MO compared to uninjected contralateral cells (Fig. 3A, A’). This was especially striking in the region of the OE where uninjected cells were maximally constricted, while the surface of adjacent morphant cells was conspicuously larger. This cellular phenotype was quantified by measuring the apical cell surface and calculating ratios between the mean size of uninjected and injected cells within the same area of individual embryos (Fig. 3B-E, B_1_-D_1_). While injection of lineage tracer alone had no effect on cell surface area (Fig. 3B, E), injection of *lrp2* MO resulted in significantly larger cell surfaces of about three times the size of uninjected constricting cells (Fig. 3C, E). Reintroduction of *lrp2* significantly ameliorated defective constriction of *lrp2* morphant cells (Fig. 3D, E). The clear distinction in size between cells of injected and uninjected side as well as in clonally distributed targeted cells (Fig. 3F) strongly indicated that Lrp2 depletion in *Xenopus* cell-autonomously impaired AC.

**Fig. 3:**
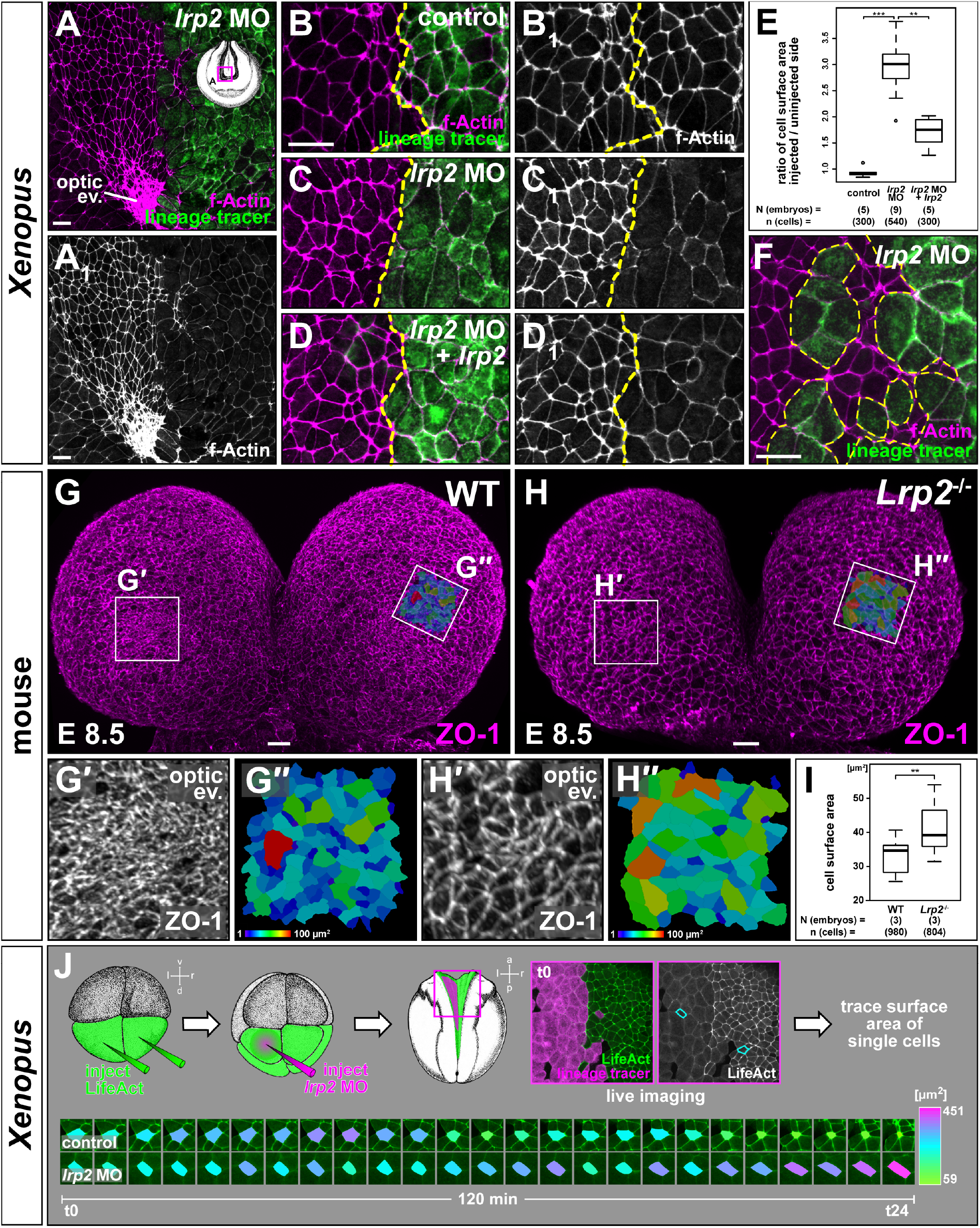
Lrp2 is cell-autonomously required for efficient apical constriction. Analysis of apical constriction (AC) in forebrain neural plate (NP) cells. (**A, A_1_**) f-Actin revealed larger apical cell surface in *lrp2* morpholino oligomer (MO)-injected cells compared to uninjected side; lack of AC in optic evagination (ev.) area on injected side. (**B-E**) Quantification of cell surface areas in unilaterally injected lineage tracer-only controls (**B**, **B_1_**), *lrp2* morphants (**C**, **C_1_**) and morphants with re-introduced *lrp2* (**D**, **D_1_**). (**E**) Cell surface area ratios calculated between injected and uninjected sides; Wilcoxon rank sum test. (**F**) Constricting NP cells intermingled with *lrp2* morphant cells demonstrated cell autonomy of AC failure. (**G, H**) Frontal views on forebrain area of wild type (WT; **G**) and *Lrp2*^−/−^ (**H**) 7 somite stage mouse embryos at embryonic day (E) 8.5.; ZO-1 delineates cell borders. (**G’**, **H’**) Magnification of optic ev. area as indicated in (**G, H**); increased cell surface in mutants. (**G”**, **H”**) Color-coded maps (areas indicated in **G, H**) visualize cell surface area. (**I**) Graphical representation of results from (**G”, H”**); four areas from each embryo were analyzed; Student’s t-test. (**J**) Live imaging (suppl. movie S2) using LifeAct; failure of AC despite sustained actin dynamics in morphants cells. Single cell surface area tracking revealed size fluctuations in all cells; final AC in control cells only. Scale bars (**A-D, F**): 25 μm; (**G, H**): 20 μm.

Consistent with these findings in *Xenopus* we also detected significantly larger cell surface areas in LRP2-deficient mouse forebrain neuroepithelial cells compared to WT stained for the tight junction protein zonula occludens-1 (ZO-1) in whole mount E8.5 embryos (Fig. 3G-I). The differences in apical surface size were especially apparent within the OE. While cells were highly constricted in the WT (Fig. 3G, G’, G”, I), mutant cells in the same area had significantly larger cell surfaces (Fig. 3H, H’, H”, I), suggesting that LRP2 deficiency also impairs AC in the mouse.

To analyze the effect of Lrp2 deficiency on the dynamic cell shape changes during AC, we applied live imaging on *Xenopus* embryos injected bilaterally with LifeAct, an *in vivo* marker for Actin dynamics, combined with unilateral *lrp2* MO injection (Fig. 3J; supp. movie 2). Quantitative analysis of cell surface size demonstrated size fluctuation over time in control cells that ultimately finalized AC. Morphant cells showed similar size fluctuations, but failed to constrict apically. However, Actin dynamics, as judged by transient protrusions and Actin movement within cells, were not affected (supp. movie 2). Together, this suggests that Lrp2 does not mediate AC by controlling Actin dynamics.

### Remodeling of apical membrane is impaired in Lrp2-deficient cells

AC leads to a decrease in apical surface area, thus creating a surplus of apical membrane. Surplus membrane arranges into structures such as ruffles, filamentous spikes / villi or spherical blebs, which can be regarded as a short-term storage of membrane prior to its resorption and redistribution by endocytic pathways (Gauthier et al., 2012). Microvilli-like membrane protrusions have been observed on apically constricting cells during gastrulation (Kurth and Hausen, 2000; Lee and Harland, 2010) and neurulation (Löfberg, 1974; Schroeder, 1970). A dynamic population of villous structures is present on epithelial cells during cellularization in *Drosophila*, and retraction of these structures by endocytosis culminates in apical cell flattening (Fabrowski et al., 2013). We thus asked whether membrane protrusions also play a role for Lrp2-mediated AC. In WT mouse embryos at E8.5 (7 somites), neuroepithelial cells mostly harbored microvilli-like filamentous protrusions in both WT controls and LRP2-deficient embryos (Fig. S3A, B). During the next hours of development, filamentous protrusions progressively receded in WT embryos and multiple bleb-like protrusions formed instead (Fig. S3C). In *Lrp2^−/−^* embryos, cells failed to retract their filamentous protrusions (Fig. S3D), reminiscent of a failure to retract villous structures in endocytosis-deficient *Drosophila* embryos (Fabrowski et al., 2013). Indeed, endocytosis was strongly impaired upon Lrp2 deficiency, as morphant cells with large apical surfaces failed to take up fluorescently labeled dextran from the medium, which was in contrast readily found intracellularly in uninjected control cells (Fig. S3E). At E9.5, we noticed strong outward bulging in cells with bigger apical diameters in *Lrp2^−/−^* embryos as compared to WT controls (Fig. S3F-H), and similar outward bulging in Lrp2-deficient *Xenopus* NP cells (Fig. S3I, I’), indicating that removal of excess apical membrane had ultimately failed. These data strongly suggest that Lrp2 as an endocytic receptor is involved in the process of eliminating surplus apical membrane, a prerequisite for efficient AC of neuroepithelial cells.

### Impaired planar cell polarity caused by loss of Lrp2 function

In addition to the impairment in AC, we had noticed widening of the caudal NP upon *lrp2* LOF in *Xenopus* (Fig. 2L, M), in agreement with LRP2-dependent caudal neurulation defects in mouse and human (Rebekah Prasoona et al., 2018; Wicher et al., 2005). Neurulation at the hindbrain and spinal cord level crucially depends on CE movements, i.e. mediolateral narrowing and concomitant anteroposterior tissue lengthening (Sutherland et al., 2020). Since these mediolateral cell intercalations are governed by the PCP pathway, we asked whether cell polarity was affected upon *lrp2* LOF. In *Xenopus*, asymmetric apical membrane localization of the core PCP component Vangl2 delineates regions undergoing CE. Already at early neurula stages (st. 13 / 14), Vangl2 localizes asymmetrically in NP cells at the hindbrain / spinal cord level (see Fig. S2 in Ossipova et al., 2015b) which starts to be prominently narrowed by CE, while it is in contrast barely detectable in the forebrain region, which remains wide and does not converge during these early stages of neurulation.

Here, we observed a regionally and subcellularly distinct distribution of pigment granules in NP cells up to mid-neurula stages (st. 15 / 16; Fig. S4A) which matched the localization of Vangl2 shown by (Ossipova et al., 2015b). While in the forebrain region, pigment was distributed symmetrically throughout individual cells (Fig. S4A, A”), it conspicuously localized in an asymmetric fashion in cells from the hindbrain level caudalward (Fig. S4A, A’). Interestingly, in *lrp2* morphants, in which the NP remained widened on the injected side (Fig. 4A, B), asymmetry at the hindbrain level was disrupted, as pigment granules distributed evenly along the cell periphery (Fig. 4B_1_, B_1’_, B_1”_), indicating that Lrp2 was required for planar polarity of NP cells at early / mid-neurula stages. Localization of Lrp2 itself shifted from the enrichment seen in constricting cells at early / mid-neurulation (st. 15; cf. Fig. 1B) to an asymmetric localization towards the medio-anterior aspect of single cells at st. 16 (Fig. 4C, C’, C’_1,2_).

**Fig. 4:**
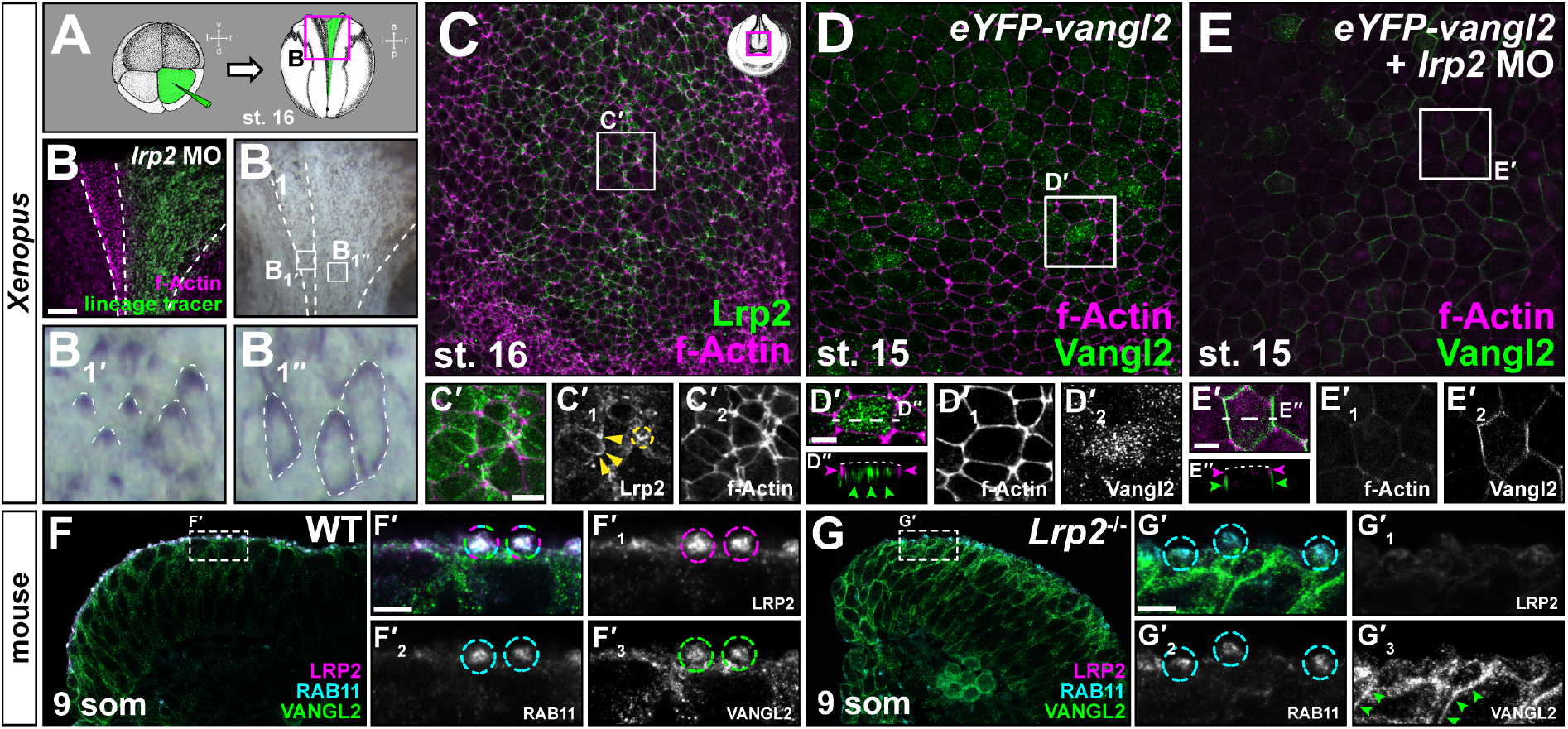
Lrp2 is required for planar cell polarity and regulates subcellular localization of Vangl2. Analysis of planar cell polarity in embryos en face (**A-E**) and on coronal forebrain sections (**F**, **G**). (**A**) Scheme for unilateral injection of morpholino oligomer (MO) targeting *lrp2;* area for analysis at stage (st.) 16 in **B** indicated by box. (**B**) Dorsal (d) view of mid- / hindbrain area; wide on injected side, f-Actin delineates cell borders. (**B_1_**) Brightfield image of same embryo shows pigment granule localization towards anterior (a) in uninjected control cells (B?); circumferential distribution in apically wide morphant cells (**B_1’_**). (**C**) Beginning of asymmetric Lrp2 distribution, f-Actin delineates cell borders. (**C’**) Magnification of area indicated in (**C**); medio-anterior distribution of Lrp2 (arrowheads in **C’_1_**), also in apically constricted cells (circle in **C’_1_**). (**C’_2_**) f-Actin single channel. (**D**, **E**) *eYFP-vangl2* injected into A1-lineage in 4-8 cell embryos, detected at st. 15 using immunofluorescence for GFP together with f-Actin staining. Dotted pattern of eYFP-Vangl2 (6 / 6 embryos; **D**, **D’**; magnification of inset in **D**), sub-apically in vesicle-like structures (**D”**; optical section indicated in **D’**). Upon *lrp2* MO injection, Vangl2 localized at cell borders (7 / 9 embryos, two independent experiments; **E**, **E’**), subapically in basolateral membrane (**E”**, optical section as indicated in **E’**). (**D’_1, 2_, E’_1, 2_**,) Single channels. (**F**, **G**) LRP2 and VANGL2 distribution at 9 somite (som) stage. (**F’_1-3_, G’_1-3_**) Magnifications as indicated by box in (**F**, **G**) and single channels thereof. Apical co-localization (dashed circles) of LRP2 and VANGL2 in RAB11-positive compartments in wild type (WT; **F**, **F’**) compared to absence of VANGL2 from RAB11-positive compartments in *Lrp2*^−/−^ cells (**G**, **G’**), arrowheads in (**G’_3_**) indicate re-localization of VANGL2 to basolateral membrane. l: left, p: posterior, r: right, v: ventral. Scale bar in (**B**): 100 μm, (**C’**, **D’**, **E’**): 10 μm, (**F’**, **G’**): 5 μm.

At mid-to late neurula stages the forebrain area narrows, leading to rapid convergence of the anterior neural folds. This suggests that planar asymmetry of PCP components plays a role in the forebrain area from mid-to late neurula stages onwards. To test whether this process is influenced by Lrp2, we first assessed the dynamics of Vangl2 localization in the forebrain area at mid-to late neurula stages. Low doses (to avoid a GOF phenotype) of *eYFP-vangl2* were injected into the neural lineage and detected using an anti-GFP antibody (Fig. 4D, Fig. S4B, C). A temporally dynamic pattern of Vangl2 subcellular localization was observed in the forebrain region. In embryos ≤ st. 16, Vangl2 was restricted to the cytoplasm and localized subapically in vesicular structures (Fig. 4D’, D’_1,2_, D”; Fig. S4B). From st. 17 onwards, Vangl2 re-distributed to the membrane (Fig. S4C) and showed a conspicuous asymmetric localization towards the medio-anterior aspect of individual cells (arrowheads in Fig. S4C, C’, C’_1,2_). Since re-localization of Lrp2 appeared slightly earlier (st. 16) than Vangl2 redistribution (st. 17), we were prompted to test whether Lrp2 is required for the localization of Vangl2 in the forebrain region. When *eYFP-Vangl2* was co-injected with *lrp2* MO (Fig. 4E) and analyzed at st. 15 (i.e. before Vangl2 re-distribution occurs in controls), we observed that Vangl2 was shifted to the lateral cell membranes and localized subapically, basal to the apical actin ring (Fig. 4E’,E’_1,2_, E”).

Consistent with this finding in *Xenopus*, LRP2 was similarly required for correct VANGL2 distribution in the mouse NP (Fig. 4F, G). In WT samples, LRP2 colocalized with VANGL2 in condensed apical structures, which were identified as recycling endosomes by RAB11 immunoreactivity (Fig. 4F, F’, F’_1-3_). While in WT controls, only very little VANGL2 was found intracellularly in vesicular structures (Fig. 4F’, F’_3_), in receptor mutant NP cells (Fig. 4G, G’, G’_1-3_), VANGL2 predominantly localized to basolateral membrane domains (arrowheads in Fig. 4G’_3_) but hardly at the apical surface.

Together, our data from mouse and frog show that 1) the core PCP protein Vangl2 was present in forebrain area NP cells in a temporally dynamic fashion, shifting its subcellular localization from apical recycling endosomes to basolateral membrane concomitant with convergence movements in the forebrain area; 2) LRP2 co-localized with VANGL2 in apical recycling endosomes and 3) Lrp2 / LRP2 was required to prevent the premature re-distribution of Vangl2 from apical recycling endosomes to basolateral membrane.

### Lrp2 interacts with intracellular adaptors to mediate cell shape changes

The endocytic pathways of transmembrane proteins are directed by intracellular adaptors. This led us to ask how Lrp2 function is mediated intracellularly. Lrp2 harbors PDZ-binding domains (PBD) within its C-terminus and binds to a diverse set of intracellular adaptor scaffold proteins in a context-dependent manner (Gotthardt et al., 2000).

Shroom3 acts as an intracellular adaptor and scaffold protein. It binds Actin, induces AC and is crucial for NP folding in both mouse and frog (Haigo et al., 2003; Hildebrand and Soriano, 1999). In the *Xenopus* NP, *shroom3* is expressed in cells engaged in AC (Haigo et al., 2003). We found that Lrp2 strongly accumulated in apically constricted hinge point cells (Fig. 5A, A_1,2_), while it was not enriched in cells in which AC had been inhibited by MO-mediated *shroom3* LOF (Fig 5A, A_1,2_). Likewise, Lrp2 strongly accumulated apically in cells of *Xenopus* blastula stage embryos, in which AC had been induced ectopically by injection of *shroom3-myc* (Fig. 5B, B_1,2_; Haigo et al., 2003), indicating that Lrp2 was recruited to sites of *shroom3*-dependent AC. We then asked whether *shroom3*-mediated AC depends on the presence of Lrp2. In cells of the animal hemisphere, *shroom3*-induced ectopic AC manifests as excessive accumulation of pigment during blastula / gastrula stages (Haigo et al., 2003). While *shroom3-myc* efficiently induced strong ectopic AC (Fig. 5C, F), loss of *lrp2* did not entirely abrogate constriction, but significantly decreased the grade of pigment accumulation (Fig. 5D, F). Re-introduction of *lrp2* rescued the MO effect (Fig. 5E, F), proving that the modulation of AC was specific to *lrp2* LOF. These data show functional interaction of the endocytic receptor Lrp2 and the constriction-inducing scaffold protein Shroom3 at the apical surface of polarized cells to facilitate efficient AC.

**Fig. 5:**
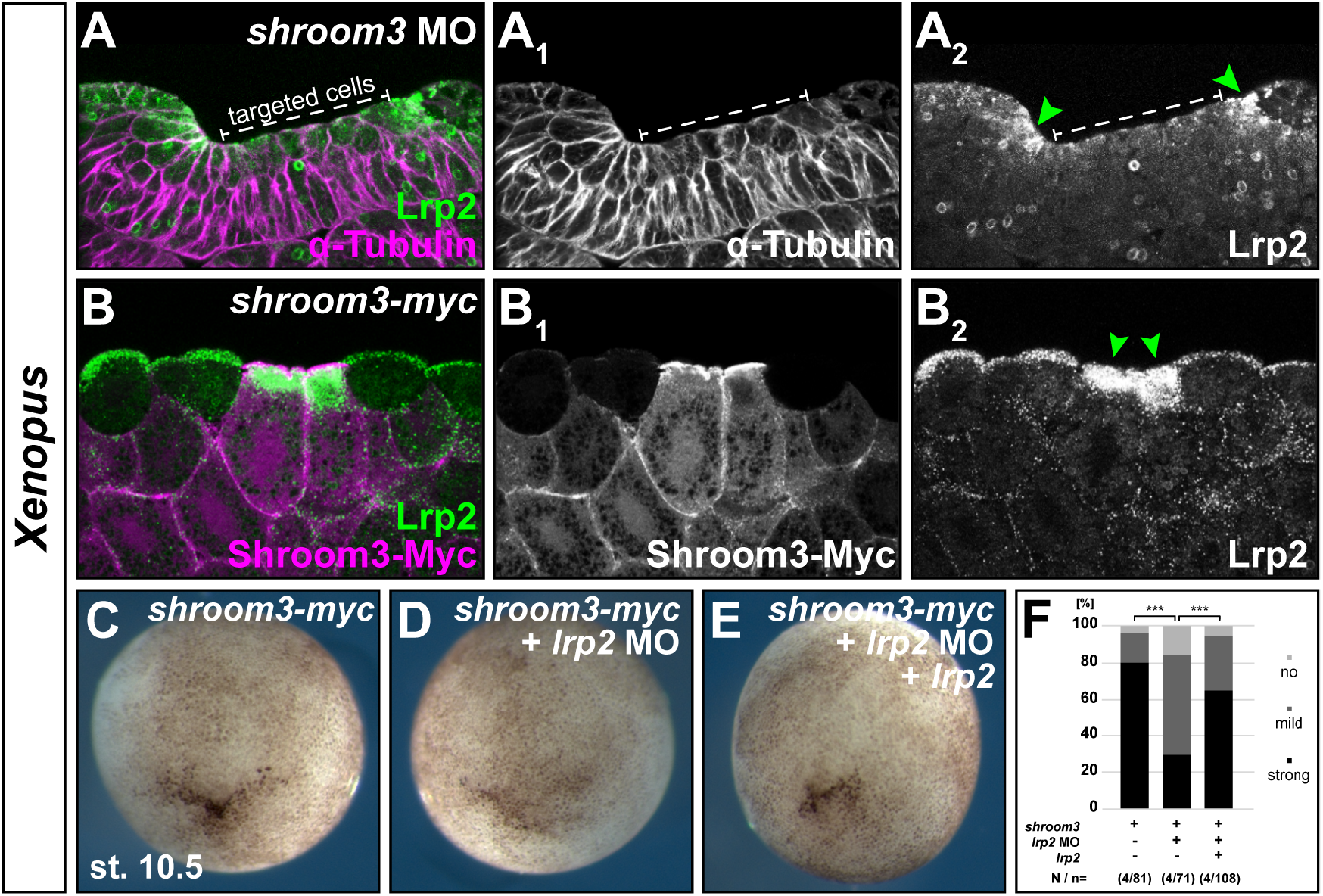
*lrp2* functionally interacts with *shroom3* in mediating apical constriction. (**A**) Transversal section through neural plate; unilateral injection of *shroom3* morpholino oligomer (MO) induced failure of apical constriction (α-tubulin illustrates wide cell surfaces; **A_1_**); apical Lrp2 accumulation in constricting hinge point cells (arrowheads in **A_2_**), lack of apical Lrp2 recruitment in targeted cells. (**B**) Section through animal cap; ectopic apical constriction induced in cells injected with *shroom3-myc* (detected by anti-Myc antibody; **B_1_**) accompanied by apical accumulation of Lrp2 (arrowheads in **B_2_**). (**C-E**) *shroom3*-induced ectopic apical constriction in animal cap cells at stage (st.) 10.5 (**C**) abrogated by co-injection of *lrp2* MO (**D**) and restored by co-injection of *lrp2*. (**F**) Quantification and statistical analysis of experiments in (**C-E**), no, mild or strong ectopic constriction was quantified. N / n = number of experiments / embryos, *χ^2^* test.

NHERF1 (Slc9a3r1) and GIPC1 are known intracellular adaptors of LRP2 (Gotthardt et al., 2000; Naccache et al., 2006; Slattery et al., 2011), however, their role in the developing neural tube has not been analyzed. NHERF1, which mediates endocytosis and trafficking of cell surface receptors, contains two PDZ domains as well as an ERM domain that enables interaction with the cytoskeleton (Weinman et al., 1998). NHERF1 overlapped with LRP2 at the apical surface of WT E8.5 mouse neural folds (Fig. S6A, A_1,2_, B, B_1,2_). Strikingly, NHERF1 was lost in *Lrp2^−/−^* mutants (Fig. S6C, C_1,2_, D, D_1,2_), suggesting a direct interaction of NHERF1 and LRP2.

Gipc1 contains one PDZ domain and is supposed to guide endocytic vesicles through the apical actin meshwork by its interaction both with receptors and myosin 6 (Aschenbrenner et al., 2003; Naccache et al., 2006). Gipc1 was localized in the NP of both mouse and *Xenopus* (Fig. 6A-C). In the mouse anterior NP at E8.5 (7 somites), GIPC1 localization showed a clear gradient: it was high in dorsolateral areas with large cell surfaces (Fig. 6A, A_1,2_, A_1’, 2’_) and lower where extensive AC occurred, such as in the midline and OE (Fig. 6A, A_1,2, A1”_, A_2”_). Similarly, in *Xenopus*, Gipc1 was absent in highly constricted cells at the border of the NP (Fig. 6B, B’), which give rise to the OE (cf. Fig. 3A). In single constricted cells located close to the NP border, Gipc1 signal was condensed subcellularly (Fig. 6C””), indicating that AC correlated with localized accumulation and disappearance of Gipc1 towards the NP border, suggestive of its degradation. Interestingly, on the tissue level, expression levels of GIPC1 and LRP2 were almost inversely correlated in the mouse. In the OE, where low levels of Gipc1 were found, Lrp2 was enriched (Fig. 1B, C; Fig. 6A_1,1”_, A_3,3”_). In agreement with the findings in mouse, in *Xenopus*, the expression levels of Gipc1 and Lrp2 did not generally correlate (Fig. 6C, compare C’_1,2_ and C”_1,2_). However, on the cellular level, co-localization was frequently found, both in large and in constricted cells (Fig. 6C”’, C”’_a_, C””, C””_a_), suggesting a spatially and temporally dynamic interaction between the two proteins.

**Fig. 6:**
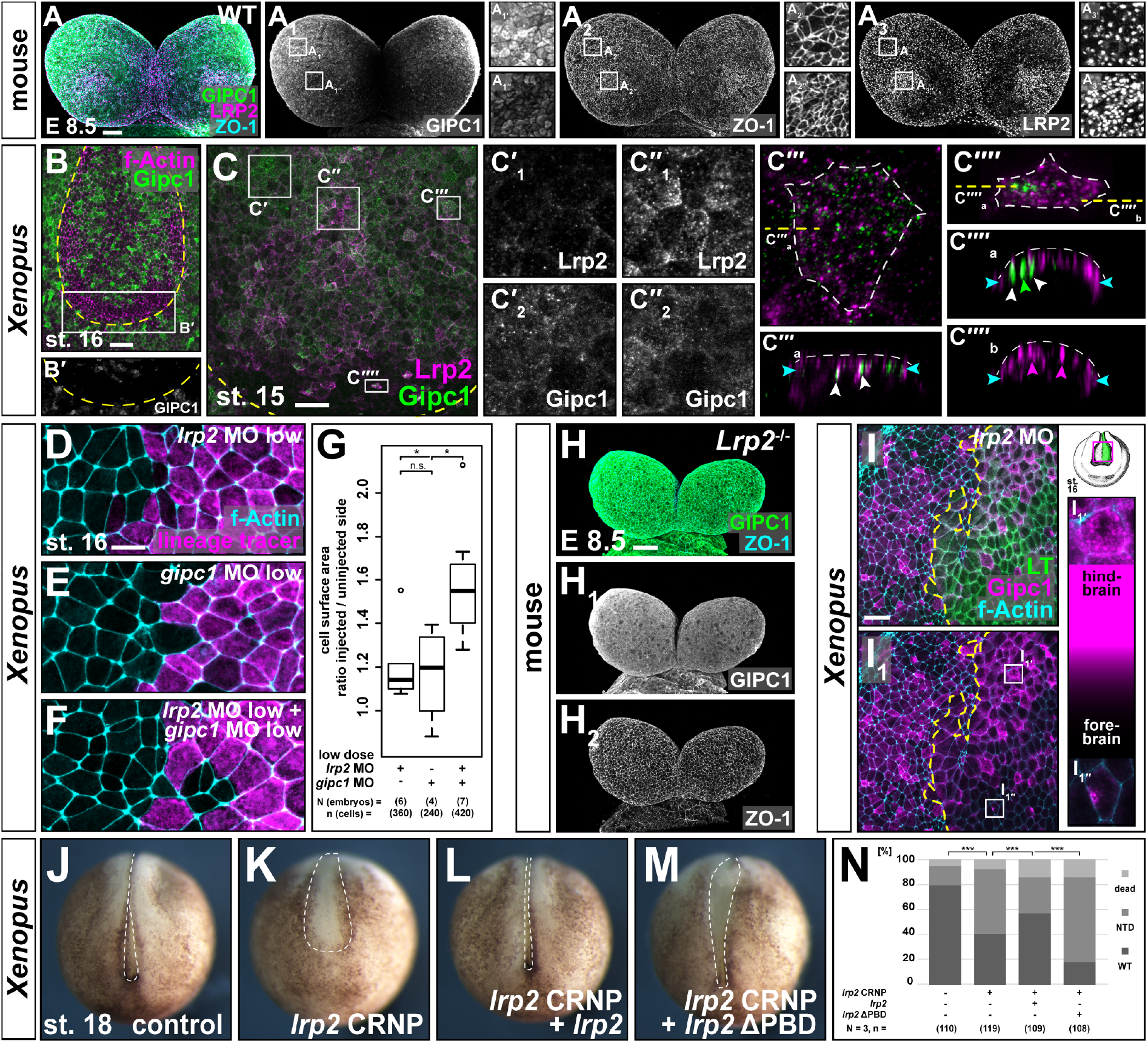
Lrp2 mediates apical constriction by functional interaction with Gipc1. (**A**) Frontal view of wild type (WT) mouse forebrain area at embryonic day (E) 8.5 (7 somites), immunofluorescence (IF) staining reveals localization of GIPC1 and LRP2, ZO-1 marks cell borders. (**A_1_-A_3_**) Single channels. (**A_1’_-A_3”_**) Magnified areas as indicated in (**A_1_-A_3_**); differential GIPC1 intensities between large and constricted cells. (**B**, **C**) Frontal views of stage (st.) 16 (**B**) and st. 15 (**C**) embryos, dashed line indicates anterior limit of neural plate; IF reveals spatially dynamic localization of Gipc1 (**B**) and spatially dynamic co-localization of Lrp2 and Gipc1 (**C**). (**B’**) Single channel of Gipc1 of magnified area indicated in (**B**). (**C’-C””**) Magnifications of boxed areas as indicated in (**C**). (**C’_1_, C”**) Single channels. Gipc1 is present in both areas with low (**C’**) and high (**C”**) amounts of Lrp2. Dispersed distribution of both Lrp2 and Gipc1 in cell with large apical surface (**C”’**), orthogonal optical section (**C””_a_**; level indicated in **C”’**) sites of Lrp2 / Gipc1 co-localization (white arrowheads). (**C””**) Cell with constricted surface; Gipc1 accumulation. (**C””_a, b_**) Both co-localization (white arrowheads) and separate localization (green / magenta arrowheads) occurs. Blue arrowheads: level of circumferential Actin belt; dashed line: apical surface in (**C”’_a_, C””_a, b_**). (**D-F**) Functional interaction of Lrp2 and Gipc1 demonstrated by individual (**D, E**) or combined (**F**) injection of low dose *lrp2 | gipc1* morpholino oligomer (MO); targeted cells with lineage tracer (LT) fluorescence. (**G**) Graphical representation of results from (**D-F**), higher than additive effect of combined MO injection; Wilcoxon rank sum test. (**H**) Frontal view of LRP2-deficient (*Lrp2^−/−^*) mouse forebrain area at E8.5 (7 somites), IF reveals localization of Gipc1, ZO-1 marks cell borders. (**H_1, 2_**) Single channels. Homogenous GIPC1 signal throughout neuroepithelium. (**I**) IF reveals mislocalization of Gipc1 in lrp2 MO-injected cells in st. 16 embryo; magnified forebrain area marked by box. Targeted cells identified by lineage tracer (LT), dashed line delineates targeted / non-targeted areas. (**I_1_**) Shown without LT. (**I_1’,1”_**) Single cells magnified as indicated in (**I_1_**); Gipc1 increases in hindbrain and decreases in forebrain area. (**J-M**) Embryos for CRISPR / Cas9 experiment all selected at the 1-cell stage and incubated until uninjected controls (**J**) reached st. 18 / 19. Injection of Cas9 ribonucleic particles (CRNP) containing sgRNA1 (**K**). (**L, M**) Co-injection of CRNP together with *lrp2* construct (**L**) or *lrp2* ΔPBD (**M**). (**N**) Graphical representation of results from (**J-M**), *χ^2^* test. Cell borders in (**D-F**, **I**) visualized by f-Actin staining. Scale bars (**A, C, H, I**): 50 μm, (**B**): 100 μm, (**D-F**): 20 μm.

To functionally analyze *gipc1*, a previously validated MO was used (Tan et al., 2001) that also induced a specific, i.e. rescuable, phenotype in the NP (Fig. S6E-I, E’-H’). Similar to the phenotype of *lrp2* morphants, MO-mediated *gipc1* LOF resulted in larger apical cell surfaces, indicating that Gipc1 also mediated AC (Fig. S6G, I). Having confirmed that both proteins were required for AC, we tested whether they functionally interacted in the process. To that end, *lrp2* or *gipc1* MOs were injected in doses low enough to induce no or only very mild AC phenotypes (Fig. 6D-G). Low dose *lrp2* or *gipc1* MO increased the median cell surface area by 14 % or 20 %, respectively (Fig. 6D, E, G), while injection of both low dose MOs led to an increase in median cell surface area of 55 % (Fig. 6F, G). The actual increase being higher than expected for an additive effect (55 % vs. 34% expected) suggested that *lrp2* and *gipc1* acted epistatically in the process of AC. To that end, *lrp2* LOF conspicuously altered the distribution of Gipc1 in both mouse and frog. In the forebrain area of *Lrp2^−/−^* mice, LRP2 deficiency completely abolished the GIPC1 expression gradient found in the WT (Fig. 6H; cf. Fig. 6A_1_), leaving all cells with a high level of GIPC1, comparable to that of GIPC1 in WT cells with a large apical surface (cf. Fig. 6A_1’_). In the forebrain area of *Xenopus lrp2* morphants, Gipc1 disappeared and only spots of asymmetrically localized Gipc1 accumulation within single cells remained (Fig. 6I, I_1_). In the hindbrain / spinal cord area, on the contrary, Gipc1 did not disappear, but was in fact upregulated (compare Fig. 6I_1’_ and 6I_1”_). Since Gipc1 has been described to bind to the C-terminal PBD of Lrp2 (Naccache et al., 2006), we asked whether this motif was required for Lrp2 function during NTC. Embryos at the 1-cell stage were injected with CRNP containing sgRNA1 (cf. Fig. S2I), which significantly induced aberrant NTC compared to uninjected controls (Fig. 6J, K, N). Co-injection of CRNP and *lrp2* (carrying the WT cytoplasmic tail) significantly reduced the amount of embryos with NTDs (Fig. 6L, N). Injecting CRNP together with *lrp2* ΔPBD, a construct lacking the last four aa which constitute the distal PBD, not only failed to rescue, but increased the number of embryos with NTDs (Fig. 6M, N). Interestingly, the NTDs in these embryos primarily manifested as widening and tissue disintegration in the caudal region of the NP.

Together, these data showed novel functional interactions of Lrp2 with the intracellular adaptor proteins Shroom3, NHERF1 and Gipc1, suggesting that spatially and temporally coordinated interaction of Lrp2 with several intracellular adaptors mediates neural morphogenesis.

## Discussion

Our functional analysis of mouse and *Xenopus* neurulation identified a conserved function of the endocytic receptor Lrp2 as a regulator of NTC. Lrp2 acted in orchestrating AC and PCP-mediated CE, two morphogenetic processes essential for proper NTC in vertebrate model organisms as well as humans (Copp et al., 2003; Wallingford et al., 2013).

### Lrp2 and its endocytic activity enable efficient apical constriction

As suggested by its accumulation in constricting cells of the NP, Lrp2 was required cell-autonomously for AC. The spatially controlled AC of neural tissue enables the creation of hinge points / hinges and the longitudinal folding of neural tissue (Colas and Schoenwolf, 2001). Impairment of AC and hinge point formation has been clearly linked to anterior NTDs (Wallingford, 2005).

How does Lrp2 as an endocytic receptor contribute to AC? Evidence is accumulating that efficient AC relies on a dual mechanism, i.e. mechanical constriction of the apical surface by actomyosin interaction accompanied by the removal of apical membrane via endocytosis (Lee and Harland, 2010; Miao et al., 2019; Ossipova et al., 2015a; Ossipova et al., 2014). In addition, AC in the NP takes place in a pulsatile manner with incremental constriction (Christodoulou and Skourides, 2015), reminiscent of a “ratchet” mechanism (Martin and Goldstein, 2014; Martin et al., 2009). During ratcheting, a phase of cell surface decrease is followed by a stabilization phase – a cyclic process, repeated until the surface is maximally constricted. It was suggested that removal of surplus membrane constitutes the mechanism leading to cell surface area stabilization (Miao et al., 2019). Our observations suggest that loss of Lrp2 does not interfere with actomyosin activation and mechanical induction of constriction. This is supported by the finding that *lrp2* LOF never entirely suppressed Shroom3-induced AC and is consistent with the lack of change in actin dynamics upon loss of Lrp2. We postulate that Lrp2 enables efficient AC by remodeling apical membrane via endocytosis to stabilize apical surface shrinkage between cycles of actomyosin constriction. This endocytic function is supported by 1) the finding that LRP2 localized to the ciliary pocket at the base of the primary cilium, a highly endocytic plasma membrane domain (Benmerah, 2013; Molla-Herman et al., 2010), 2) the severe inhibition of dextran endocytosis upon *lrp2* LOF and 3) the formation of membrane protrusions which were not removed from the constricting surface in a timely manner.

### Lrp2 controls the timing of PCP protein localization

It has been shown that endocytic uptake, recycling, intracellular re-localization and endocytic removal of uncomplexed PCP proteins at cell junctions are essential processes to establish PCP-mediated cell and tissue polarization (Eaton and Martin-Belmonte, 2014). Acquisition of cell polarity brought about by endocytic trafficking is thus a temporally dynamic process, illustrated e.g. by Vangl2 localization. In zebrafish dorsal mesoderm, Vangl2 is first detected cytoplasmically in vesicle-like structures and relocates to the membrane shortly prior to the onset of PCP-dependent cell polarization (Roszko et al., 2015). It accumulates asymmetrically just before initiation of CE. The temporal differences in WT Vangl2 localization prior to and during convergence of the *Xenopus* neural folds that were observed here match the dynamics in zebrafish mesoderm. This timeline suggests that the finalization of forebrain NTC requires a temporal control of PCP. A requirement for endocytosis in this process is underlined by our finding that loss of Lrp2 induced a premature and aberrant re-localization of Vangl2 from apical cytoplasmic compartments (Rab11-positive recycling endosomes) to basolateral membrane in mouse and frog. Interestingly, despite a localization to the basolateral membrane, Vangl2 did not accumulate in a planar polarized fashion. This suggests that the correct succession of events during PCP-dependent neural fold convergence requires Lrp2-mediated endocytic trafficking 1) for the temporal restriction of Vangl2 to cytoplasmic vesicular compartments and 2) to control the levels and asymmetric distribution of PCP proteins in the subapical / basal membrane. The latter is in line with the phenotype of PCP protein GOF, which also disrupts convergence movements and cell polarity (Darken et al., 2002; Goto and Keller, 2002; Wallingford et al., 2000). We conclude that Lrp2-mediated endocytosis and trafficking is required for the precise control of timing, amount and localization of PCP proteins to drive anterior neurulation.

### Lrp2 as a hub to orchestrate AC and PCP?

A functional link between AC and PCP, especially during neural tube closure, has become increasingly evident (McGreevy et al., 2015; Nishimura et al., 2012; Ossipova et al., 2014; Ossipova et al., 2015b). Since we found that both morphogenetic processes were affected by *lrp2* LOF, Lrp2-mediated endocytosis and intracellular trafficking could be a common denominator for AC and PCP. How would an interaction between AC and PCP – each with a conspicuously different cellular outcome – be mediated by a single receptor? A likely explanation lies in the ability of Lrp2 to differentially interact with intracellular adaptors and scaffold proteins via its C-terminal cytoplasmic motifs.

Shroom3 is an intracellular adaptor and scaffold protein. We found that Lrp2 was recruited apically upon *shroom3* injection and was required for efficient Shroom3-induced AC. It remains to be tested whether a scaffold containing Lrp2 and Shroom3 is necessary for initiation of endocytosis, intracellular trafficking and efficient AC. Protein scaffolds also serve as integrators for signaling pathways and for compartmentalization of components that contribute to different pathways at different times (Pawson and Scott, 2010). A Lrp2-based scaffold might thus serve as a platform for temporospatial integration of AC and PCP (Fig. 7). Both Lrp2 and Shroom3 feature a conserved PBD (X(S/T)X(V/L)) for class | PDZ domains in their distal C-terminus. While the Shroom3 PBD remains to be functionally analyzed (Haigo et al., 2003; Hildebrand and Soriano, 1999; Lee et al., 2007), the Lrp2 PBD is functionally relevant for interaction with the class | PDZ domain-containing adaptor protein Gipc1 (Naccache et al., 2006). Since Gipc1 also dimerizes (Reed et al., 2005), it serves as a connector and may create an AC-mediating scaffold containing Lrp2, Shroom3 and Gipc1. Indeed, we demonstrate here a novel requirement for Gipc1 in facilitating AC and NTC. Lrp2-Gipc1 functional interaction is supported by the LOF phenocopy of *lrp2* and *gipc1*, their epistatic relation and the requirement for Lrp2’s PBD in rescue experiments. The influence of Lrp2 on Gipc1 localization further suggests not only a functional, but also a physical interaction between these proteins. We could thus envision a complex of Shroom3, Lrp2 and Gipc1 that facilitates efficient AC via engagement of actomyosin (Shroom3) and endocytic elimination of membrane (Lrp2). Indeed, Gipc1 dimers interact with Lrp2 and at the same time mediate the formation of a complex with Myosin6, which facilitates the trafficking of endocytic vesicles through the apical actin network (Aschenbrenner et al., 2003). Thus, Gipc1-mediated guidance of Lrp2-positive endocytic vesicles through the apical actin meshwork could account for efficient removal of apical membrane upon Shroom3-induced AC in anterior NP cells.

**Fig. 7:**
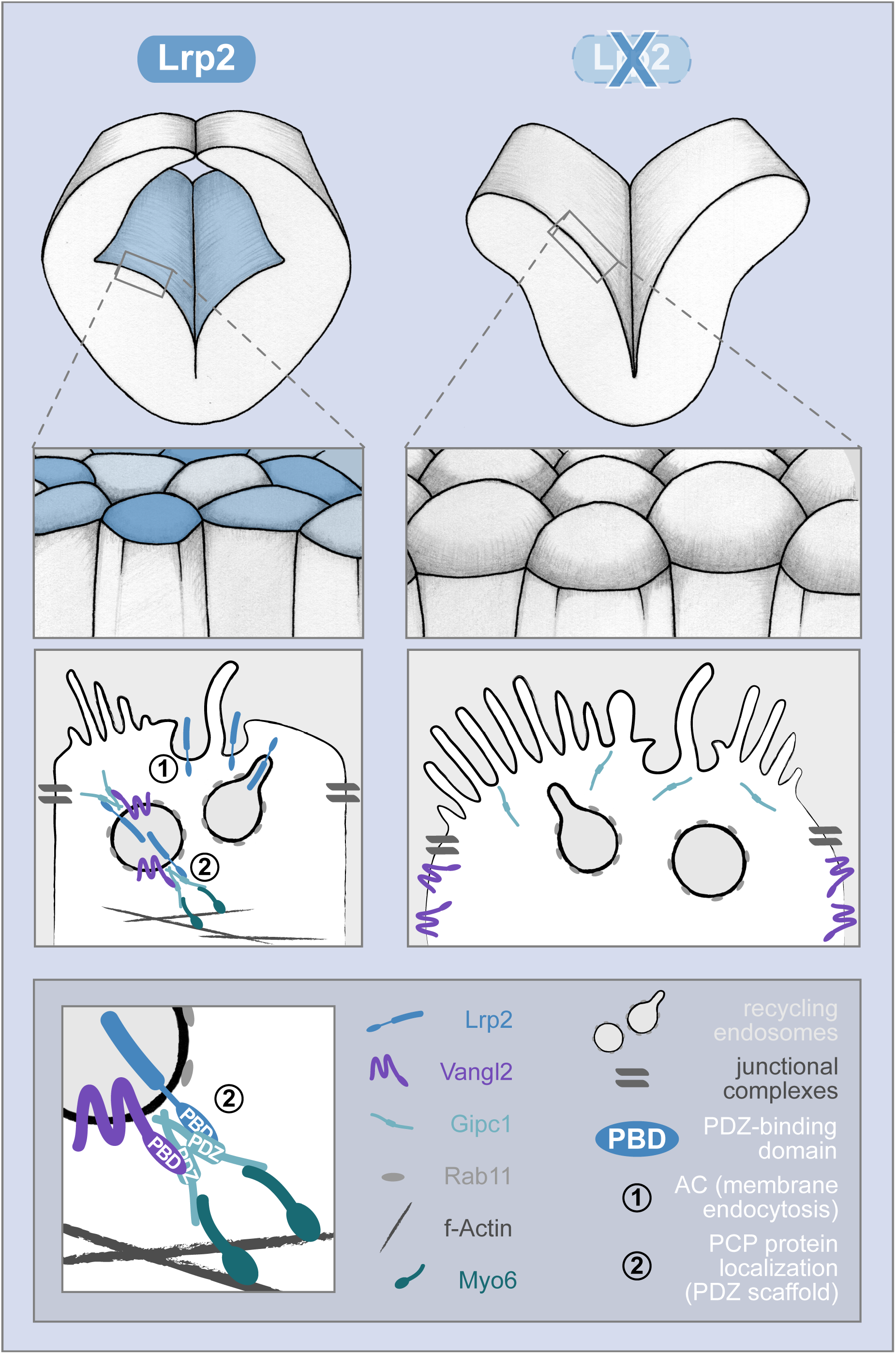
Hypothetical model of Lrp2 functional interactions in neural tube closure. The neural tube closes, facilitated by apical constriction of Lrp2-positive neuroepithelial cells (left column). Lrp2 mediates endocytic removal of apical membrane (1) as well as correct temporospatial localization of Vangl2 (2) via recycling endosomes. Lrp2 / Vangl2 interaction is likely facilitated by PDZ / PBD-mediated intracellular scaffolding via dimerized Gipc1, connecting to Myo6 and the actin cytoskeleton. Lack of Lrp2 (right column) entails defective neural tube closure due to impaired apical constriction. Removal of apical membrane fails and proper subcellular sorting of Vangl2 and Gipc1 is disturbed, leading to their mislocalization.

Our observations here suggest that a phase of Shroom3-initiated AC during early neurulation is succeeded by PCP-mediated CE to finalize anterior NTC. Such a temporal succession of AC and PCP could also be mediated through scaffolding by Gipc1 (Fig. 7). *gipc1* is clearly also involved in the establishment of PCP. It directs the localization of the core PCP protein Vangl2 and its LOF elicits PCP phenotypes (Giese et al., 2012). Vangl2 in turn is known to localize differentially in phases of AC vs. PCP. It is recruited to the apical membrane in constricting cells *in vivo* and upon ectopically induced AC (Ossipova et al., 2014; Ossipova et al., 2015a). Our temporal analysis of Vangl2 localization in the anterior NP of *Xenopus* revealed that the apical vesicular localization of Vangl2 coincided with AC, while membrane localization occurred later during neural fold convergence. This temporal pattern and the premature membrane mislocalization of Vangl2 upon *lrp2* LOF suggests that Lrp2 is required to retain Vangl2 in recycling endosomes during AC. Its controlled release from recycling endosomes would enable its re-localization to the membrane and the initiation of PCP / CE. Since Gipc1 can bind to both Lrp2 and Vangl2 it could act as an adaptor between both proteins and mediate the controlled release of Vangl2 from the recycling route. We propose that AC and PCP / CE in the anterior NP are temporally and spatially separated processes, the succession of which is regulated by Lrp2 / Gipc1-mediated endocytosis and intracellular trafficking.

### Conservation of Lrp2 function in disease etiology

Most of the clinical characteristics seen in patients with *LRP2* gene mutations are also present in the LRP2-deficient mouse models (Cases et al., 2015; Hammes et al., 2005; Kur et al., 2014; Sabatino et al., 2017; Spoelgen et al., 2005; Wicher and Aldskogius, 2008). Our report demonstrates that *Xenopus* is a new valuable model for the functional analysis of Lrp2 deficiency. In addition to the matching neural localization and LOF phenotypes between mouse and frog, *Xenopus lrp2* is expressed in DBS disease manifestation sites such as otic vesicle and pronephros (Fig. S1 and Christensen et al., 2008). This suggests that the frog will also be highly valuable for studies on human *LRP2*-related congenital disorders of organs other than the neural tube. Of note, the intracellular adaptors addressed here indeed share similar LOF phenotypes such as kidney insufficiency (Shroom3, Khalili et al., 2016; NHERF1 Shenolikar et al., 2002; Gipc1, Naccache et al., 2006) and hearing loss (NHERF1, Girotto et al., 2019; Gipc1, Giese et al., 2012 and discussion therein).

Together, our data suggest a novel role for LRP2 in the functional interaction with subapical scaffolds that are essential for proper neuroepithelial morphogenesis and neural tube closure.

Our findings here support the notion that the function of Lrp2 in these processes is conserved also in humans. Thus, combining the power of the *Xenopus* and mouse embryological models should prove highly valuable to study the mechanistic origins of human NTDs and other congenital disorders related to LRP2 dysfunction.

## Materials and methods

### Animals

#### Mouse

Animal experiments were performed according to institutional guidelines following approval by local authorities (X9005/12). Mice were housed in a 12 hour light-dark cycle with *ad libitum* food and water. The generation of mice with targeted disruption of the *Lrp2* gene on a C57BL/6NCrl background has been described (Willnow et al., 1996). Analyses of the embryonic neural tube defects were carried out in LRP2-deficient and in somite-matched wild type and heterozygous littermates on a C57BL/6NCrl background.

#### Xenopus

All animals were treated according to the German regulations and laws for care and handling of research animals, and experimental manipulations were approved by the Regional Government Stuttgart, Germany (“Vorhaben Xenopus Embryonen in der Forschung” V340/17 ZO and V349/18 ZO).

### Cloning of expression constructs

The constructs here referred to as *lrp2* and *lrp2*ΔPBD (lacking the last four amino acids which consitute the PDZ-binding domain, PBD) were generated from HA-Meg4, encoding an extracellularly truncated human megalin / LRP2 (kindly provided by Maria Paz Marzolo). HA-Meg4 contains an HA-tag N-terminally, followed by the fourth ligand binding domain as well as the transmembrane and intracellular domains. Using a standard PCR-based approach with primers containing restriction sites, the constructs were amplified and subcloned into the pCS2+ vector.

### Microinjections of morpholino oligomers and mRNA in *Xenopus*

Drop size was calibrated to 4 nl per injection and dextran tetramethylrhodamine or dextran Alexa Fluor 488 (MW 10,000, 0.5 – 1 μg / μl, Thermo Fisher Scientific) were added as a lineage tracer. Morpholino oligomers (MOs; Gene Tools) used were: *lrp2* MO (ATG-spanning, translation-blocking; 5’ AGCTCCCATCTCTGTCTCCTGC 3’), *gipc1* MO (5’UTR-located, translation-blocking; 5’ CCACGGACAGCAAATCTCACACAG 3’; Tan et al., 2001), both 0.5 – 1 pmol per injection; for epistasis experiments, 0.3 pmol MO was used each. The *gipc1, shroom3-myc* and *eYFP-vangl2* constructs contained the ORFs of *Xenopus laevis gipc1.L, shroom3.L* and *vangl2.S*, respectively. Capped mRNAs were synthesized using mMessage mMachine (Ambion), amounts per injection were: *gipc1*, 10 – 30 pg; *lrp2*, 400 pg; *lrp2*ΔPBD, 400 pg; *shroom3*, 200 pg; *eYFPvangl2*, 100 pg; *LifeAct*, 400 pg. In all experiments care was taken to exclude specimens that were not targeted correctly, i.e. in which fluorescence was not restricted to the neural plate or in which fluorescence could not be evaluated optimally at mid-neurula stages.

### CRISPR/Cas9-mediated genome editing in *Xenopus*

Single guide RNAs (sgRNAs) were designed using CRISPRscan (CRISPRscan.org; (Moreno-Mateos et al., 2015), synthesized from double-stranded template DNA using the MEGAshortscipt T7 transcription kit (Invitrogen, AM1354) and purified using the MEGAclear transcription clean-up kit (Invitrogen, AM1908). Cas9 (PNA Bio, CP01-50) ribonucleic particles (CRNPs) were assembled by heating sgRNA to 70°C followed by immediate chilling to prevent formation of secondary structures and subsequent incubation with Cas9 at 37°C for 5 min. Per injection, a volume of 8 nl containing 1 ng Cas9 / 300 pg sgRNA was delivered into the animal pole of oocytes approx. 20 min after fertilization. To evaluate editing, DNA from a pool of ten embryos was harvested by lysis at the desired stage, PCR amplicons containing the cutting site were sequenced and knock-out efficiency was calculated using the Synthego ICE online tool (ice.synthego.com; Hsiau et al., 2019).

### Dextran uptake assay

Embryos at the 8-cell stage were injected unilaterally with *lrp2* MO and raised in 0.1 x MBSH until st. 14, when the vitelline membrane was removed. Embryos were incubated in 0.1 x MBSH containing 10 ng / μl dextran tetramethylrhodamine (Invitrogen, D1817) until stage 18. They were then transferred to fresh 0.1 x MBSH and further reared until st. 20, fixed in PFA, washed, bisected at the level of the forebrain and further processed for staining and imaging.

### Whole mount *in situ* hybridization

*Xenopus* embryos were fixed for 2 h in 1 x MEMFA at room temperature and further processed for *in situ* hybridization following standard protocols (Harland, 1991). The probe for *lrp2*.L was kindly provided by André Brändli (Christensen et al., 2008).

### Immunofluorescence (IF)

#### Mouse

For standard whole mount imaging, E8.5 mouse embryos were dissected and the rostral neural plate was collected. Tissue was fixed for 1 h in 4% PFA at room temperature, washed in 1 x PBS and either dehydrated in a methanol series to be stored in 100% MetOH at −20 °C or directly subjected to the IF protocol. Embryos were permeabilized with PBS-Triton X-100 (0.1 %) for 15 min at room temperature and blocked with this solution containing 1 % donkey serum and 2 % BSA for 6 h at room temperature. Incubation with primary antibodies was carried out for 48 h at 4 °C using the following dilutions: sheep anti-LRP2 antiserum (1:5000), kindly provided by the laboratory of Renata Kozyraki, rabbit anti-LRP2 (Abcam cat. # ab76969; 1:1000), mouse anti-ZO-1 (Invitrogen cat. # 33-9100; 1:100), mouse anti-ARL13b (UC Davis / NIH, NeuroMab cat. # 75-287; 1:500), rabbit anti-GIPC1 (Alomone cat. # APZ-045; 1:200). Bound primary antibodies were visualized using secondary antibodies conjugated with Alexa Fluor 488, 555, 647 after overnight incubation (Abcam, 1:500). All tissues were counterstained with DAPI (Invitrogen, cat. # 62248). Embryos were mounted with ProLong Gold Antifade Mountant (Invitrogen, cat. # P36934) in between two cover slips, using Secure-Seal Spacer (Invitrogen, cat. # S24737).

For IF on cryosections, PFA-fixed embryos were infiltrated with 15 % and 30 % sucrose in PBS up to 1 h, embedded in O.C.T. (Tissue-Tek^®^ O.C.T., Sakura Finetek, cat. # sa-4583) and cut into 10 μm coronal sections. Standard IF staining was carried out by incubation of tissue sections with primary antibodies overnight at 4 °C at the following dilutions: mouse anti-acetylated tubulin (Sigma cat. # T7451; 1:1000), mouse anti-RAB11 (BD Transduction Laboratories cat. # 610657; 1:200), rabbit anti-VANGL2 (1:500) kindly provided by the laboratory of Mireille Montcouquiol, sheep anti-LRP2 antiserum (1:5000), kindly provided by the laboratory of Renata Kozyraki, rabbit anti-NHERF1 (Alomone, cat. # APZ-006; 1:500). Bound primary antibodies were visualized using secondary antibodies conjugated with Alexa Fluor 488, 555, 647 after 1 h incubation at room temperature (Abcam, 1:500). All tissues were counterstained with DAPI (Invitrogen, cat. # 62248). Sections were mounted with Dako fluorescence mounting medium (Agilent, cat. # S302380-2).

For whole mount STED imaging, E9.5 mouse embryos were collected and the neural tube was slit open using insect needles along the dorsal midline from caudal to rostral. The neural folds were precisely cut above the heart and placed on a sterile filter (Millipore, cat. # MCEP06H48) within a drop of PBS (1 x) in a petri dish. The floor plate at the level of the cephalic flexure was pinched in order to unfold the tissue with the ventricular part facing up. Filters were placed in a 6-well plate containing DMEM / 10 % FCS and explants were incubated at 37°C, with 5 % CO_2_ and 95 % humidity for 3-4 hours to flatten and recover. The explants were washed gently in 1 x PBS, fixed for 1 h in 4 % PFA, and subjected to the standard IF protocol described above. Highly cross-absorbed secondary antibodies Alexa Fluor Plus 594 (Invitrogen, cat. # A32744) and Atto 647N (Active Motif, cat. # 15038) were used. Explants were flat-mounted in ProLong Gold Antifade Mountant (Invitrogen, cat. # P36934) to obtain optimal resolution.

#### Xenopus

Embryos were fixed in a solution of 4 % PFA in 1 x PBS for 1 h at room temperature or overnight at 4° C, then washed in 1 x PBS to remove fixative. For whole-mount staining, the vitelline membrane was carefully removed and embryos were transferred to CAS blocking reagent (Invitrogen, cat. # 008120). For staining of sections, embryos were fixed in 1 x MEMFA, embedded in 2% agarose and sectioned on a VIBRATOME series 1000. The following primary antibodies were used: rabbit anti-Lrp2 (Abcam cat. # ab76969), mouse anti-MYC (clone 9E10, Abcam cat. # ab32), monoclonal mouse anti-α-Tubulin (clone DM1A, Sigma cat. # T6199), goat anti-Gipc1 (Sigma-Aldrich, cat. # SAB2500463), chicken anti-GFP (Invitrogen, cat. # A10262). Where possible, subtype-specific secondary antibodies coupled to either AlexaFluor 488 or 555 were used (Invitrogen). AlexaFluor 405, 488 or 555-coupled phalloidin (Invitrogen) was used to stain filamentous actin. DNA was stained with Hoechst 33342 (Invitrogen) to visualize nuclei.

### Microscopy

#### Confocal microscopy, image processing and analysis

Image acquisitions of mouse tissue sections and mouse neural folds were carried out using a Leica SP8 confocal microscope with either HC Pl Apo CS2 63× NA 1.4 oil immersion objective for sections or HC Pl Apo 20× NA 0.75 MultiIMM with glycerol immersion for whole mount imaging. The raw data from whole mount mouse embryos were acquired close to the Nyquist sampling limit with a z-piezo stepper (80 nm pixel size, 0.5 μm z-step size 12 bit, dynamic range). In all samples, Alexa Fluor 488 was excited by a 488 nm laser, detection at 500 – 550 nm, Alexa Fluor 555 was excited by a 555 nm laser, detection at 570 – 620 nm, Alexa Fluor 647 was excited by a 633 nm or 647 nm laser, detection at 660 – 730 nm and DAPI was excited at 405 nm, detection at 420 – 450 nm with a pinhole set to 1 AU. All samples that were compared either for qualitative or quantitative analysis were imaged under identical settings for laser power, detector and pixel size.

Confocal Z-stacks of whole mount neural folds were subjected to a background correction and processed by deconvolution with the CMLE algorithm and theoretical PSF in order to obtain an improved signal to noise ratio and axial and spatial resolution using Huygens Professional software (Scientific Volume Imaging). The deconvolution was applied to all image sets prior further segmentation and analysis steps with the Imaris Software.

*Xenopus* samples were imaged on a Zeiss LSM5 Pascal or LSM700 confocal microscope.

#### STED imaging

En face STED images of mouse cephalic explants were taken with a Leica SP8 TCS STED microscope (Leica Microsystems) equipped with a pulsed white-light excitation laser (WLL; ~80 ps pulse width, 80 MHz repetition rate; NKT Photonics) and two STED lasers for depletion at 592 nm and 775 nm. The system was controlled by the Leica LAS X software. Dual-color STED imaging was performed by sequential excitation of Alexa Fluor Plus 594 at 590 nm and Atto 647N at 647 nm. For emission depletion the 775 nm STED laser was used. Time-gated detection was set from 0.3 – 6 ns. Two hybrid detectors (HyD) were used at appropriate spectral regions separated from the STED laser to detect the fluorescence signals. The emission filter was set to 600 – 640 nm for Alexa Fluor Plus 594 and to 657 – 750 nm for Atto 647N. Images were sequentially acquired with a HC PL APO CS2 100 x / 1.40 NA oil immersion objective (Leica Microsystems), a scanning format of 1,024 × 1,024 pixels, 8-bit sampling, 16 x line averaging and 6 x optical zoom, yielding a voxel dimension of 18.9 x 18.9 nm. In addition to every STED image, a confocal image with the same settings but just 1 x line averaging was acquired.

#### Scanning electron microscopy

E8.5 embryos were dissected and fixed in 0.1 M sodium cacodylate buffer (pH 7.3 / 7.4) containing 2.5 % glutaraldehyde. Rinsing in cacodylate buffer was followed by a postfixation step in 2 % OsO_4_ for two hours. Samples were dehydrated in a graded ethanol series, osmicated, dried in critical point apparatus (Polaron 3000), coated with gold / palladium MED 020 (BAL-TEC) and examined using a Zeiss scanning electron microscope Gemini DSM 982.

#### Transmission electron microscopy

After dissection, mouse embryos at E9.5 were fixed with 3 % formaldehyde in 0.2 M HEPES buffer, pH 7.4, for 30 min at RT followed by postfixation with 6 % formaldehyde / 0.1 % glutaraldehyde in 0.2 M HEPES buffer for 24 h at 4 °C. Samples were stained with 1 % OsO_4_ for 2 h, dehydrated in a graded ethanol series and propylene oxide and embedded in Poly/Bed^R^ 812 (Polysciences, Inc., Eppelheim, Germany). Ultrathin sections were contrasted with uranyl acetate and lead citrate. Sections were examined with a Thermo Fisher Morgagni electron microscope, digital images were taken with a Morada CCD camera and the iTEM software (EMSIS GmbH, Münster, Germany). The same software was used to manually measure the size of the average cell diameter.

#### Video-documentation of neural development

For videography of actin dynamics, *LifeAct* mRNA was injected into both dorsal blastomeres of albino embryos at the four-cell stage, followed by unilateral injection of *lrp2* MO at the eight-cell stage. Correct targeting was verified at early neural plate stages and only embryos targeted correctly into the neural lineage were used. A time series of single plane confocal images (pinhole > 1 Airy unit to increase optical section thickness, one frame per minute) was recorded on a Zeiss LSM 700 using a 20 x objective.

For brightfield imaging of neurulation, timelapse sequences of embryos injected unilaterally with *lrp2* MO were recorded at two frames per minute from stage 13 onwards on a Zeiss stereomicroscope (SteREO Discovery.V12) with an AxioCam HRc (Zeiss).

### Measurements and statistics

#### Neural plate width quantification

For neural plate width measurement, control and treated embryos were photographed frontally after *in situ* hybridization for the pan-neural marker *sox3* and analyzed in ImageJ. The floor plate, easily identified by the lightest staining along the rostrocaudal neural midline, was marked and the widest part of the anterior neural plate was measured orthogonally to the midline. For each embryo, the ratio between injected and uninjected side was calculated.

#### Cell surface area quantification

The anterior neural folds of matching somite stage wild type and mutant mouse embryos were subjected to cell surface area analysis. Regions of interest of the same size were chosen (4 per sample) and cropped in 3D in IMARIS (Bitplane) from the whole mount images. Using a maximum intensity projection in Fiji they were transformed into 2D data sets. The Imaris Cell segmentation module was used for analysis, excluding incomplete cells from the edges. Manual adjustment was performed if necessary and final cell surface area parameters were extracted. The complete dataset was subjected to statistical analysis using an unpaired T-test.

In *Xenopus*, cell surface areas were measured manually using ImageJ. To that end, at least 30 cells from corresponding areas of uninjected and injected side of each embryo (i.e. 60 cells per embryo) were analyzed. The mean surface area of each side was calculated and used to determine a ratio between injected and uninjected side.

### Statistical analysis

Statistical tests used to analyze the data were done using Prism 7 software (GraphPad) or Statistical R and are mentioned in the respective figure legends. Significance was scored as follows: *p* ≥ 0.05: not significant; *p* < 0.05: *; *p* < 0.01: **; *p* < 0.001: ***. p-value levels, numbers of specimens and biological replicates are reported in the figures or figure legends.

## Acknowledgements

We thank Nora Mecklenburg for her intellectual input to the project. The professional assistance of Anje Sporbert and Matthias Richter with confocal microscopy, of Mrs. Schrade, Bettina Purfürst and Christina Schiel with electron microscopy and of Martin Lehmann and Hannes Gonschior with STED microscopy is gratefully acknowledged. Many thanks to Mireille Montcouquiol for kindly providing the VANGL2 antibody. We appreciate Manfred Ströhmann’s work in the mouse husbandry and Anke Scheer’s technical assistance. We thank Gary Lewin for critical reading of the manuscript and Thomas Willnow for acquisition of financial support for the project. Many thanks to Martin Blum and members of the zoology department for support and discussion, Tim Ott for advice on CRISPR analysis and Ann-Kathrin Burkhart and Niklas Schaedler for their work on Lrp2 during the initial steps of this project.

## Data availability

n/a

