## Supplemental Figures for "Neural tube closure requires the endocytic receptor Lrp2 and its functional interaction with intracellular scaffolds"

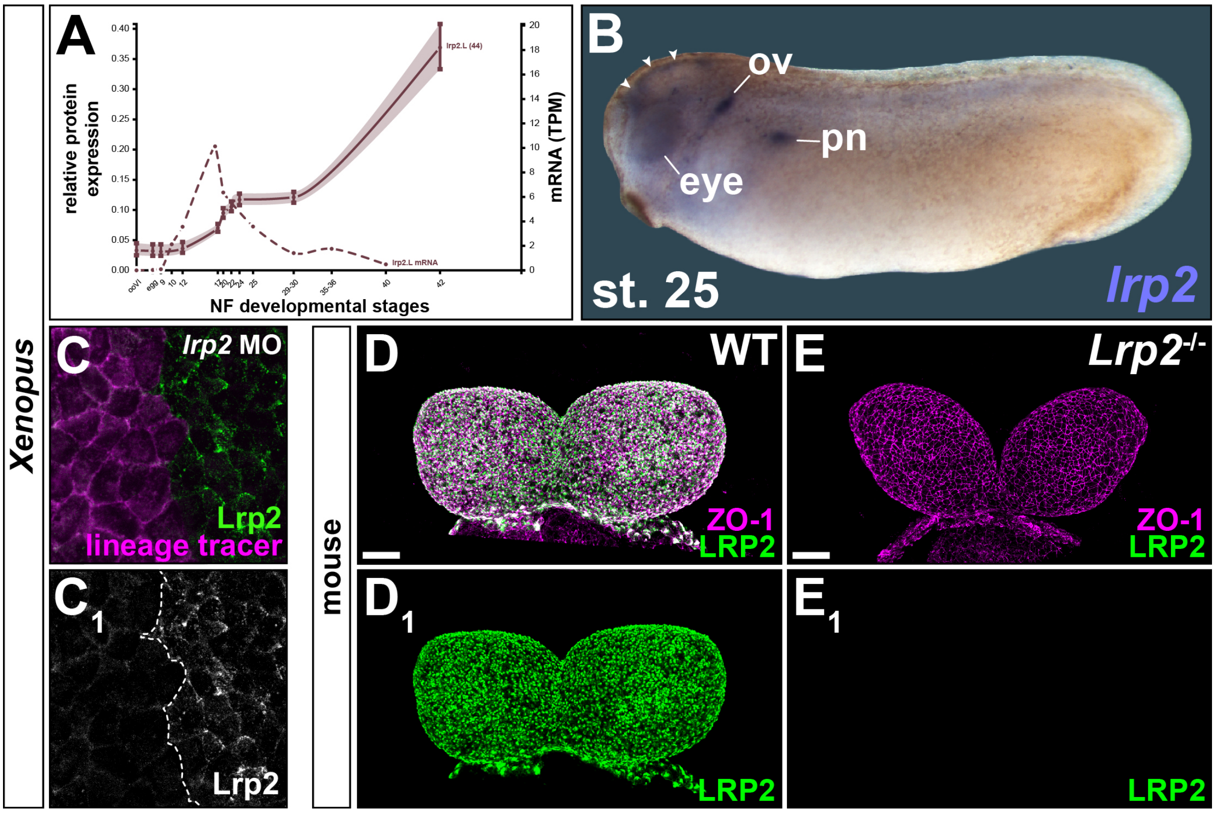


**Fig. S1: Lrp2 expression and protein depletion by knock-down in *Xenopus* and knock-out in mouse**. (**A**) mRNA and protein expression profile of *Xenopus lrp2*.L (from Xenbase.org). (**B**) *Xenopus lrp2* expressed in brain (arrowheads), eye anlage, otic vesicle (ov) and proximal pronephros (pn) in stage (st.) 25 tailbud embryo. (**C**) Morpholino oligomer (MO) decreases Lrp2 expression in injected cells; single channel (**$C_1$**) for clarity. (**D, E**) Frontal views of anterior neural folds of wild type (WT; **D**) and Lrp2$^{-/-}$ (**E**) mouse embryos at embryonic day (E) 8.5. (**$D_1$**, **$E_1$**) WT LRP2 expression (**$D_1$**) lost in Lrp2$^{-/-}$ (**$E_1$**). ZO-1 labels cell boundaries. Scale bars (**D**, **E**): 50 µm.

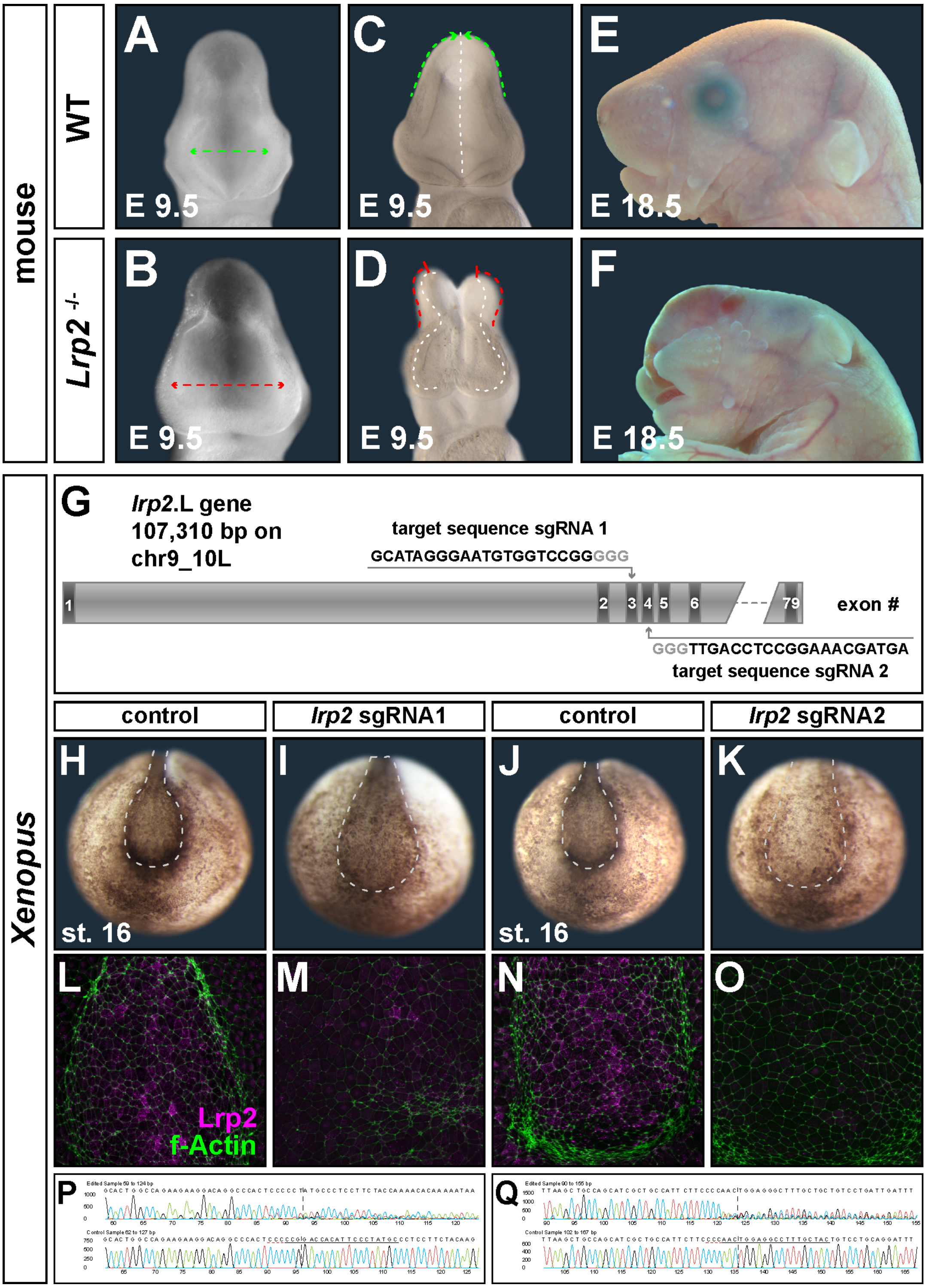


**Fig. S2: Mouse phenotypes and CRISPR-Cas9-mediated genome editing of *lrp2*.L in *Xenopus laevis*.** (**A**-**F**) Additional forebrain phenotypes of *Lrp2*$^{-/-}$ mice. (**A-D**) Heads of mice, frontal views, at embryonic day (E) 9.5. Normally wide (dotted line in **A**) vs. dilated dorsal forebrain (dotted line in **B**) in 65% (22 / 34) of *Lrp2*$^{-/-}$ embryos. (**C**) Green dotted lines indicate closed neural tube (NT) in WT. (**D**) Failure of NT closure (red dotted lines) in 35% (12 / 34) of *Lrp2*$^{-/-}$ embryos. (**E, F**) Anencephaly as a consequence of impaired NT closure in *Lrp2*$^{-/-}$ E18.5 embryos. (**G**) Schematic representation of *Xenopus laevis lrp2*.L gene structure; target sequence for single guide (sg) RNA-binding in exon 3 and 4 is shown, protospacer-adjacent motif (PAM) in gray letters. (**H-O**) Injection of Cas9-ribonucleic particles (CRNP) assembled with sgRNA1 or 2 into zygotes impaired neural plate narrowing and lengthening (**H-K**) and reduced Lrp2 (**L-O**) in CRISPants compared to controls. f-Actin marks cell circumference. (**P**, **Q**) Analysis of sequence flanking target sites confirmed editing in CRNP-injected samples (top row) compared to control samples (lower row).

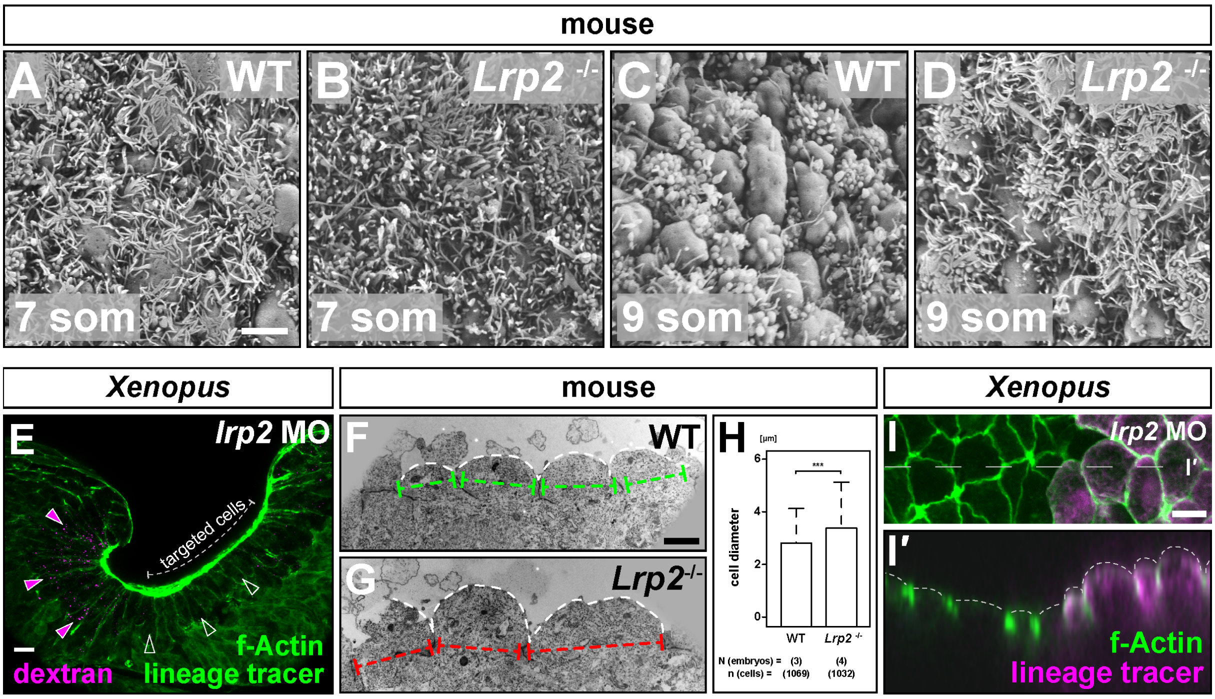


**Fig. S3: No efficient remodeling of apical surface in cells with apical constriction failure.** (**A**-**D**) Scanning electron micrographs of mouse neuroepithelial cells at embryonic day (E) 8.5. Note prominent filamentous, microvilli-like protrusions on cells of wild type (WT; **A**) and *Lrp2*-deficient (*Lrp2*$^{-/-}$; **B**) embryos at the 7 somite (som) stage. In 9 som embryos, filamentous protrusions are reduced in the WT (**C**), but persist in *Lrp2*$^{-/-}$ (**D**). (**E**) Transverse section of *Xenopus* forebrain area at stage (st.) 19; cells targeted by *lrp2* morpholino oligomer (MO) identified by enlarged apical surface and cytosolic lineage tracer fluorescence. Embryos were incubated in medium containing fluorescently coupled dextran from st. 14 - 19. Note dextran signal in constricted cells (filled arrowheads) but absence of dextran from MO-targeted cells (empty arrowheads). F-actin staining indicates cell borders. (**F**, **G**) Transmission electron micrographs of E 9.5 coronal ultrathin sections revealed cells with normal apical cell diameters (green lines) and moderate bulging in the WT (**F**) compared to increased cell diameter (red lines) and excessive bulging in *Lrp2*$^{-/-}$ neural plate cells (**G**). (**H**) Quantification and statistical analysis of cell diameters. Student's t-test. (**I**) En face view of neural plate, apically enlarged *lrp2* morpholino oligomer (MO)-targeted cells (lineage tracer$^{+}$; f-Actin marks cell borders) bulged outward as seen in orthogonal view (**I′**; level of optical section indicated in **I**). Scale bars (A-D): 2 µm; (E): 20 µm; (F, G): 2 µm; (I): 10 µm.

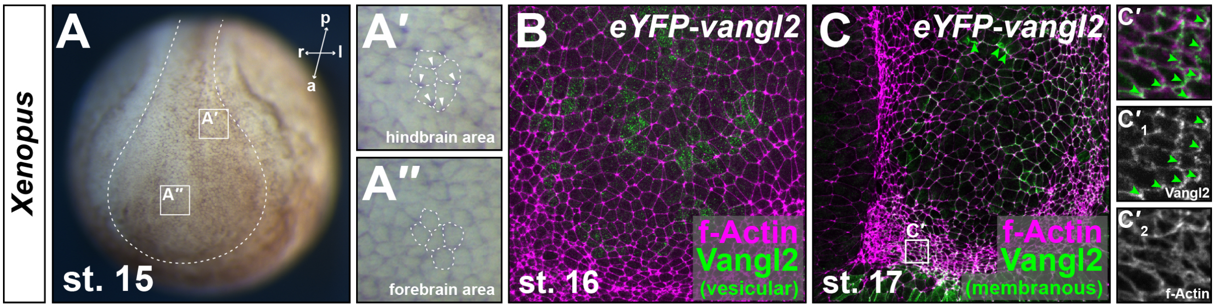


**Fig. S4: Dynamic planar cell polarity in the *Xenopus* neural plate.** (**A**) Brightfield image; frontal view of a stage (st.) 15 embryo, note asymmetric pigment accumulation in anterior (a) aspect of cells (outlined by dashed line) in the hindbrain area (arrowheads in **A'**) compared to symmetric circumferential distribution in forebrain area cells (**A''**). (**B**, **C**) *eYFP-vangl2* was injected into A1-lineage in 4-8 cell embryos and detected using indirect immunofluorescence for GFP, f-Actin visualizes cell borders. eYFP-Vangl2 localized in dotted, vesicle-like pattern up to st. 16 (**B**; 6 embryos, st. 14-16); re-distribution to asymmetric membrane localization from st. 17 onwards (arrowheads in **C** and **C'**; 8 embryos, st. 17 / 18). Magnified area (**C'**) as indicated in (**C**); (**C'$_{1,2}$**) single channels shown in grayscale. l: left, p: posterior, r: right.

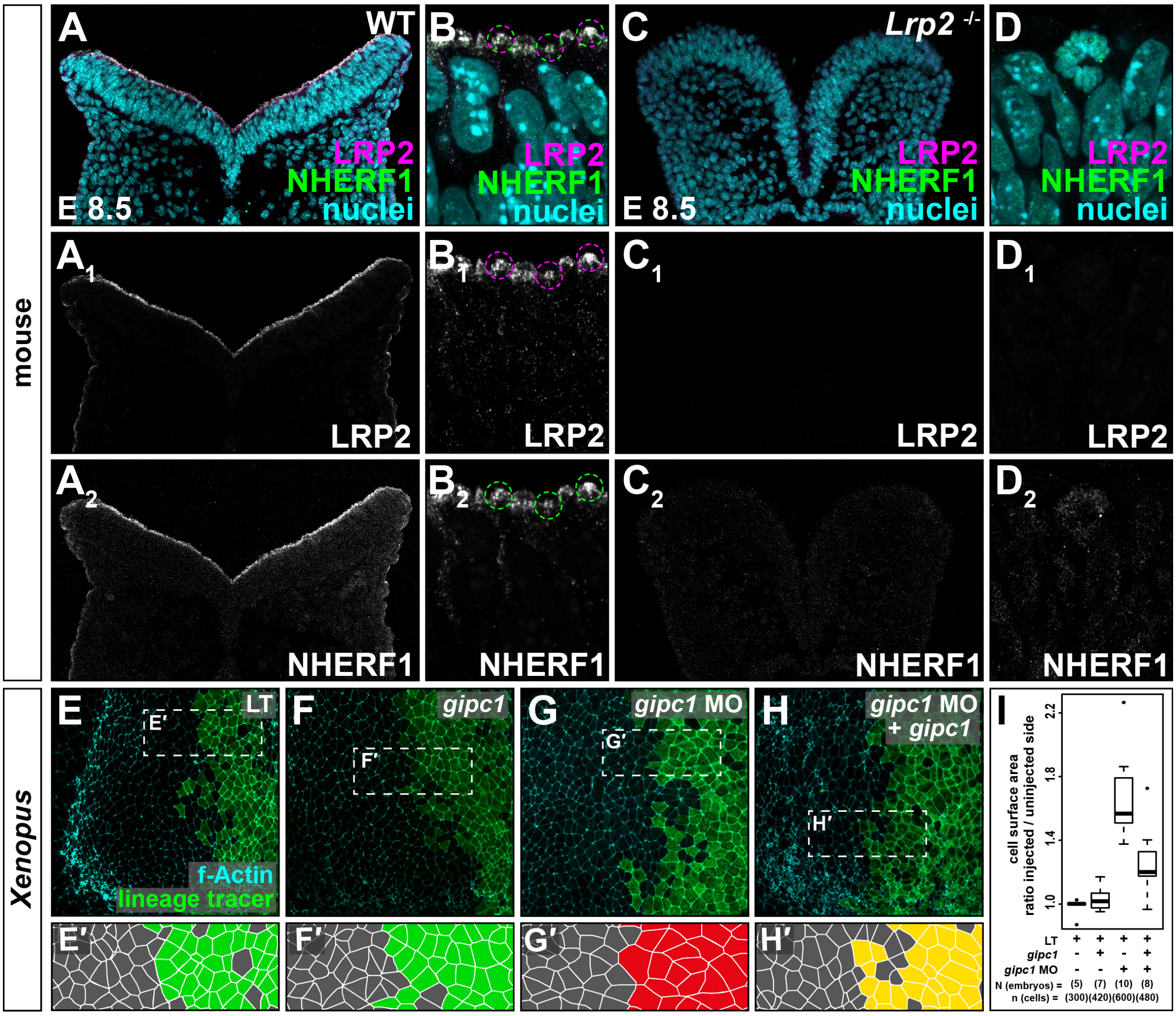


**Fig. S6: Lrp2-interacting scaffold proteins in neural tube closure.** (**A-D**) Immunofluorescence reveals co-localization of LRP2 and NHERF1 on coronal sections of wild type (WT; **A**, **B**) and loss of both markers in LRP2-deficient (*Lrp2*$^{-/-}$; **C**, **D**) mice at embryonic day (E) 8.5 (9 somites). (**A**$_{1,2}$ **- D**$_{1,2}$) Channels shown in grayscale. (**E-I**) Morpholino oligomer (MO)-mediated *gipc1* loss-of-function (LOF) phenocopies the Lrp2 LOF phenotype. 4-8 cell embryos were injected into one dorsal animal blastomere with lineage tracer (LT) and MO / mRNA as indicated; cell borders delineated by f-Actin staining using fluorescently labeled phalloidin. (**E'-H'**) Cell outlines from boxes indicated in (**E-H**), colors represent severity of phenotype. No impairment of constriction upon injection of LT (**E**, **E'**) or *gipc1* (**F**, **F'**; green), severe impairment in MO-injected cells (**G**, **G'**; red), amelioration of impaired constriction upon re-introduction of *gipc1* in MO-injected cells (**H**, **H'**; yellow). (**I**) Graphical representation of experimental data from (**A-D**), Wilcoxon rank sum test.

**Supplemental movie 1:** ***lrp2*** **is required for neural tube closure in** ***Xenopus***

**Supplemental movie 2:** ***lrp2*** **is required for apical constriction, but dispensable for actin dynamics in the neural plate**
